# Deep Neural Networks and Kernel Regression Achieve Comparable Accuracies for Functional Connectivity Prediction of Behavior and Demographics

**DOI:** 10.1101/473603

**Authors:** Tong He, Ru Kong, Avram J. Holmes, Minh Nguyen, Mert R. Sabuncu, Simon B. Eickhoff, Danilo Bzdok, Jiashi Feng, B.T. Thomas Yeo

## Abstract

There is significant interest in the development and application of deep neural networks (DNNs) to neuroimaging data. A growing literature suggests that DNNs outperform their classical counterparts in a variety of neuroimaging applications, yet there are few direct comparisons of relative utility. Here, we compared the performance of three DNN architectures and a classical machine learning algorithm (kernel regression) in predicting individual phenotypes from whole-brain resting-state functional connectivity (RSFC) patterns. One of the DNNs was a generic fully-connected feedforward neural network, while the other two DNNs were recently published approaches specifically designed to exploit the structure of connectome data. By using a combined sample of almost 10,000 participants from the Human Connectome Project (HCP) and UK Biobank, we showed that the three DNNs and kernel regression achieved similar performance across a wide range of behavioral and demographic measures. Furthermore, the generic feedforward neural network exhibited similar performance to the two state-of-the-art connectome-specific DNNs. When predicting fluid intelligence in the UK Biobank, performance of all algorithms dramatically improved when sample size increased from 100 to 1000 subjects. Improvement was smaller, but still significant, when sample size increased from 1000 to 5000 subjects. Importantly, kernel regression was competitive across all sample sizes. Overall, our study suggests that kernel regression is as effective as DNNs for RSFC-based behavioral prediction, while incurring significantly lower computational costs. Therefore, kernel regression might serve as a useful baseline algorithm for future studies.

## 1. Introduction

Deep neural networks (DNNs) have enjoyed tremendous success in machine learning (Lecun et al., 2015). As such, there has been significant interest in the application of DNNs to neuroscience research. DNNs have been applied to neuroscience in at least two main ways. First, deep learning models have been used to simulate actual brain mechanisms, such as in vision (Khaligh-Razavi and Kriegeskorte, 2014; Yamins et al., 2014; Eickenberg et al., 2017) and auditory perception (Kell et al., 2018). Second, DNNs have been applied as tools to analyze neuroscience data, including lesion and tumor segmentation (Pinto et al., 2016; Havaei et al., 2017; Kamnitsas et al., 2017b; G. Zhao et al., 2018), anatomical segmentation (Wachinger et al., 2018; X. Zhao et al., 2018), image modality/quality transfer (Bahrami et al., 2016; Nie et al., 2017; Blumberg et al., 2018), image registration (Yang et al., 2017; Dalca et al., 2018), as well as behavioral and disease prediction (Plis et al., 2014; van der Burgh et al., 2017; Vieira et al., 2017; Nguyen et al., 2018).

Deep neural networks can perform well in certain scenarios and tasks, where large quantities of data are unavailable, e.g., winning multiple MICCAI predictive modeling challenges involving image segmentation (Choi et al., 2016, Kamnitsas et al., 2017a, Hongwei Li et al., 2018). Yet, the conventional wisdom is that DNNs perform especially well when applied to well-powered samples, for instance, the 14 million images in ImageNet (Russakovsky et al., 2015) and Google 1 Billion Word Corpus (Chelba et al., 2014). However, in many neuroimaging applications, the available data often only involve hundreds or thousands of participants, while the associated feature dimensions can be significantly larger, such as entries of connectivity matrices with upwards of 100,000 edges. Consequently, we hypothesize that in certain neuroimaging applications, DNNs might not be the optimal choice for a machine learning problem (Bzdok and Yeo, 2017). Here, we investigated whether DNNs can outperform classical machine learning for behavioral prediction using resting-state functional connectivity (RSFC).

RSFC measures the synchrony of resting-state functional magnetic resonance image (rs-fMRI) signals between brain regions (Biswal et al., 1995; Fox and Raichle, 2007; Buckner et al., 2013), while participants are lying at rest without any explicit task. RSFC has been widely used for exploring human brain organization and mental disorders (Smith et al., 2009; Assaf et al., 2010; Power et al., 2011; Yeo et al., 2011; Bertolero et al., 2017). For a given brain parcellation scheme (e.g., Shen et al., 2013; Gordon et al., 2016; Glasser et al., 2017; Eickhoff et al., 2018), the parcels can be used as regions of interest (ROIs), such that a whole brain (or cortical) RSFC matrix can be computed for each participant. Each entry of the RSFC matrix corresponds to the strength of functional connectivity between two brain regions. In recent years, one of the most influential developments in neuroimaging has been the use of the RSFC matrices for predicting the attributes (e.g., age or fluid intelligence) of individual participants (Dosenbach et al., 2010; Finn et al., 2015; Smith et al., 2015; Rosenberg et al., 2016; Dubois et al., 2018; Reinen et al., 2018; Weis et al., 2019). Consequently, there have been many studies developing new techniques to improve RSFC-based behavioral prediction (Amico and Goñi, 2018; Nostro et al., 2018; Parisot et al., 2018; Kashyap et al., 2019; Yoo et al., 2019).

In this work, we compared kernel regression with three DNN architectures in RSFC-based behavioral prediction. Kernel regression is a non-parametric classical machine learning algorithm (Murphy, 2012) that has previously been utilized in various neuroimaging prediction problems, including RSFC-based behavioral prediction (Raz et al., 2017; Zhu et al., 2017; Kong et al., 2019; Li et al., 2019). Our three DNN implementations included a generic, fully-connected feedforward neural network, and two state-of-the-art DNNs specifically developed for RSFC-based prediction (Kawahara et al., 2017; Parisot et al., 2017, 2018). An initial version of this study utilizing only the fluid intelligence measure in the HCP dataset has been previously presented at a workshop (He et al., 2018). By using RSFC data from nearly 10,000 participants and a broad range of behavioral (and demographic) measures from the HCP (Smith et al., 2013; Van Essen et al., 2013) and UK Biobank (Sudlow et al., 2015; Miller et al., 2016), this current extended study represents one of the largest empirical evaluations of DNN’s utility in RSFC-based fingerprinting.

## 2. Methods

### 2.1 Datasets

Two datasets were considered: the Human Connectome Project (HCP) S1200 release (Van Essen et al., 2013) and the UK Biobank (Sudlow et al., 2015; Miller et al., 2016). Both datasets contained multiple types of neuroimaging data, including structural MRI, rs-fMRI, and multiple behavioral and demographic measures for each subject.

HCP S1200 release comprised 1206 healthy young adults (age 22-35). There were 1,094 subjects with both structural MRI and rs-fMRI. Both structural MRI and rs-fMRI were acquired on a customized Siemens 3T “Connectome Skyra” scanner at Washington University at St. Louis. The structural MRI was 0.7mm isotropic. The rs-fMRI was 2mm isotropic with TR of 0.72s and 1200 frames per run (14.4 minutes). Each subject had two sessions of rs-fMRI, and each session contained two rs-fMRI runs. A number of behavioral measures were also collected by HCP. More details can be found elsewhere (Van Essen et al., 2012; Barch et al., 2013; Smith et al., 2013).

The UK Biobank is a prospective epidemiological study that has recruited 500,000 adults (age 40-69) between 2006-2010 (Sudlow et al., 2015). 100,000 of these 500,000 participants will be brought back for multimodal imaging by 2022 (Miller et al., 2016). Here we considered an initial release of 10065 subjects with both structural MRI and rs-fMRI data. Both structural MRI and rs-fMRI were acquired on harmonized Siemens 3T Skyra scanners at three UK Biobank imaging centres (Cheadle Manchester, Newcastle, and Reading). The structural MRI was 1.0mm isotropic. The rs-fMRI was 2.4mm isotropic with TR of 0.735s and 490 frames per run (6 minutes). Each subject had one rs-fMRI run. A number of behavioral measures were also collected by the UK Biobank. More details can be found elsewhere (Elliott and Peakman, 2008; Sudlow et al., 2015; Miller et al., 2016; Alfaro-Almagro et al., 2018).

### 2.2 Preprocessing and RSFC

We utilized ICA-FIX MSM-All grayordinate rs-fMRI data provided by the HCP S1200 release (HCP S1200 manual; Van Essen et al., 2012, 2013; Glasser et al., 2013; Smith et al., 2013; Griffanti et al., 2014; Salimi-Khorshidi et al., 2014). To eliminate residual motion and respiratory-related artifacts (Burgess et al., 2016), we performed further censoring and nuisance regression (Kong et al., 2019; Li et al., 2019) Runs with more than 50% censored frames were discarded (Pruett et al., 2015; Gordon et al., 2016; Smyser et al., 2016; Kong et al., 2019; Li et al., 2019). Figure S1 shows the distribution of the number of uncensored frames across subjects.

Consistent with previous studies from our group (Kebets et al., 2019; Li et al., 2019), we considered 400 cortical (Schaefer et al., 2018) and 19 sub-cortical (Fischl et al., 2002; Glasser et al., 2013) ROIs to ensure whole-brain coverage. The preprocessed rs-fMRI time courses were averaged across all grayordinate locations within each ROI. RSFC was then computed using Pearson’s correlation of the averaged time courses for each run of each subject (with the censored frames excluded for the computation). The RSFC was averaged across all runs, resulting in one 419 x 419 RSFC matrix for each subject.

In the case of the UK Biobank, we utilized the 55 x 55 RSFC (Pearson’s correlation) matrices provided by the Biobank (Miller et al., 2016; Alfaro-Almagro et al., 2018). The 55 ROIs were obtained from a 100-component whole-brain spatial-ICA (Beckmann and Smith, 2004), of which 45 components were considered to be artifactual (Miller et al., 2016).

### 2.3 FC-based prediction setup

We considered 58 behavioral measures across cognition, emotion and personality from the HCP (Table S1; Kong et al., 2019). By restricting the dataset to participants with at least one run (that survived censoring) and all 58 behavioral measures, we were left with 953 subjects. 23, 67, 62 and 801 subjects had 1, 2, 3 and 4 runs respectively.

**Table 1.**
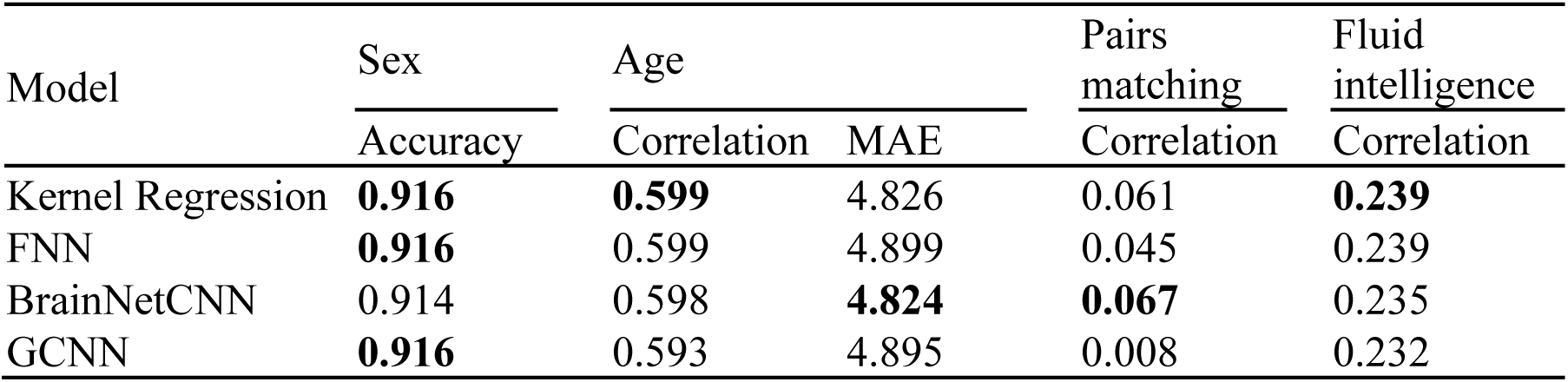
Prediction performance of four behavioral and demographic measures in the UK Biobank. For age (MAE), lower values imply better performance. For all the other measures, larger values imply better performance. **Bold** indicates the best performance, although it does not imply statistical significance. There was no statistical difference between kernel regression and the DNNs for all behavioral and demographic measures after correcting for multiple comparisons (q < 0.05). MAE refers to mean absolute error. Correlation refers to Pearson’s correlation. We note that simply predicting the median age in the training set would have yielded an MAE of 6.194.

In the case of the UK Biobank, we considered four behavioral and demographic measures: age, sex, fluid intelligence and pairs matching^1^ (number of incorrect matches). By restricting the dataset to participants with 55 x 55 RSFC matrices and all four measures, we were left with 8868 subjects.

For both datasets, kernel regression and three DNNs were applied to predict the behavioral and demographic measures of individual subjects based on individuals’ RSFC matrices. More specifically, the RSFC data of each participant was summarized as an N x N matrix, where N is the number of brain ROIs. Each entry in the RSFC matrix represented the strength of functional connectivity between two ROIs. The entries of the RSFC matrix were then used as features to predict behavioral and demographic measures in individual participants.

### 2.4 Kernel ridge regression

Kernel regression (Murphy, 2012) is a non-parametric classical machine learning algorithm. Let *y* be the behavioral measure (e.g., fluid intelligence) and *c* be the RSFC matrix of a test subject. Let *y_i_* be the behavioral measure (e.g., fluid intelligence) and *c_i_* be the RSFC matrix of the *i*-th training subject. Roughly speaking, kernel regression will predict the test subject’s behavioral measure to be the weighted average of the behavioral measures of all training subjects: *y* ≈ *∑_iɛtraining set_ Similarity*(*c_i_*, *c*)*y_i_*, where *Similarity*(*c_i_*, *c*) is the similarity between the RSFC matrices of the test subject and *i*-th training subject. Here, we simply set *Similarity*(*c_i_*, *c*) to be the Pearson’s correlation between the lower triangular entries of matrices *c_i_* and *c*, which is effectively a linear kernel. In practice, an *l*_2_ regularization term is needed to avoid overfitting (i.e., kernel ridge regression). The level of *l*_2_ regularization is controlled by the hyperparameter λ. More details are found in Appendix A1.

### 2.5 Fully-connected neural network (FNN)

Fully-connected neural networks (FNNs) belong to a generic class of feedforward neural networks (Lecun et al., 2015) illustrated in Figure 1. An FNN takes in vector data as an input and outputs a vector. An FNN consists of several fully connected layers. Each fully connected layer consists of multiple nodes. Data enters the FNN via the input layer nodes. Each node (except input layer nodes) is connected to all nodes in the previous layer. The values at each node is the weighted sum of node values from the previous layer. The weights are the trainable parameters in FNN. The outputs of the hidden layer nodes typically go through a nonlinear activation function, e.g., Rectified Linear Units (ReLU; *f*(*x*) = *max*(0,*x*)), while the output layer tends to be linear. The value at each output layer node typically represents a predicted quantity. Thus, FNNs (and neural networks in general) allow the prediction of multiple quantities simultaneously. In this work, the inputs to the FNN are the vectorized RSFC (i.e., lower triangular entries of the RSFC matrices) and the outputs are the behavioral or demographic variables we seek to predict.

**Figure 1.**
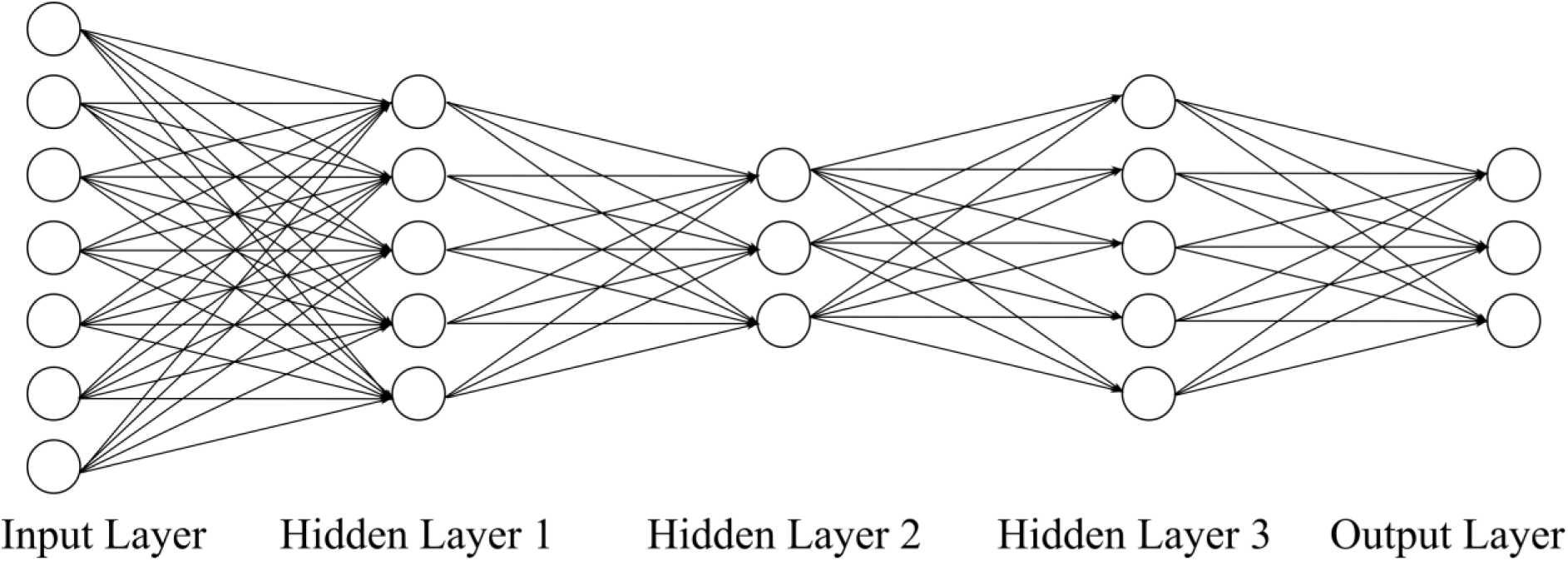
Schematic of a feedforward neural network (FNN). An FNN takes in vectorized RSFC matrix entries as inputs and outputs behavioral or demographic predictions. An FNN consists of an input layer, several hidden layers (three layers are shown here) and an output layer. The number of nodes in the input layer is equal to the number of elements in the lower triangular portion of the RSFC matrix. The number of nodes in the output layer is typically equal to the number of behavioral measures we are predicting. The number of hidden layers and number of nodes in the hidden layers are among the many hyperparameters that have to be tuned.

### 2.6 BrainNetCNN

One potential weakness of the FNN is that it does not exploit the (mathematical and neurobiological) structure of the RSFC matrix, e.g., RSFC matrix is symmetric, positive definite and represents a network. On the other hand, BrainNetCNN (Kawahara et al., 2017) is a specially designed DNN for connectivity data, illustrated in Figure 2. BrainNetCNN allows the application of convolution to connectivity data, resulting in significantly less trainable parameters than the FNN. This leads to less parameters, which should theoretically improve the ease of training and reduce overfitting issues. In this work, the input to the BrainNetCNN is the *N* × *N* RSFC matrix and the outputs are the behavioral or demographic variables we seek to predict.

**Figure 2.**
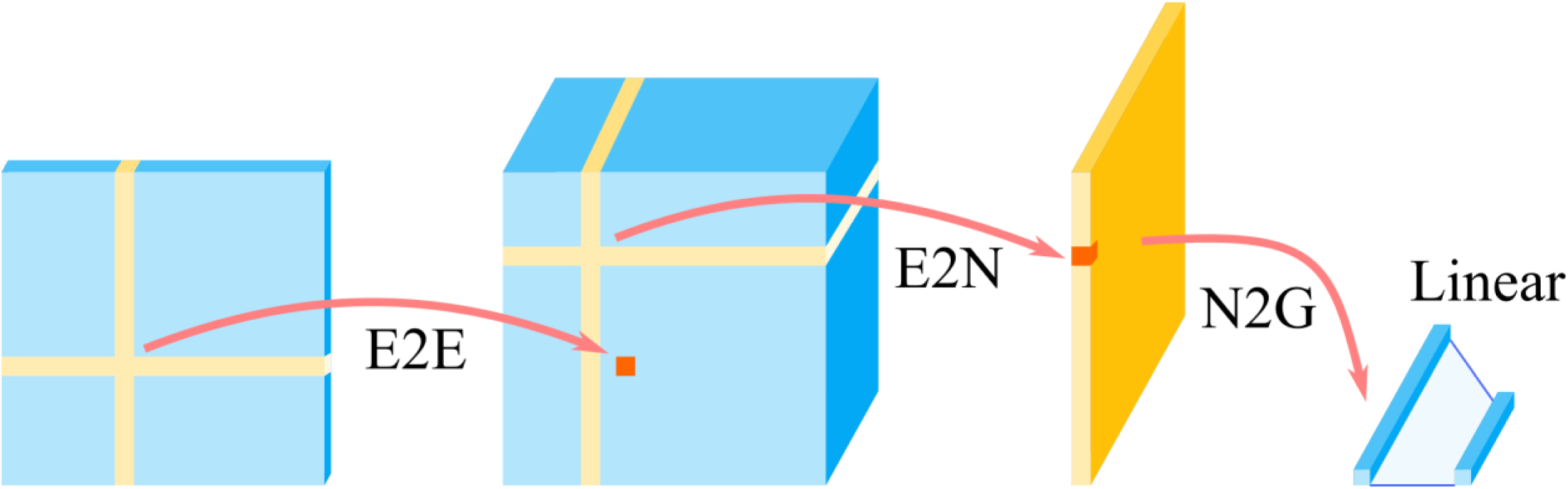
Schematic of the BrainNetCNN. (Kawahara et al., 2017). The BrainNetCNN takes in the RSFC matrix as an input and outputs behavioral or demographic predictions. BrainNetCNN consists of four types of layers, Edge-to-Edge (E2E) layer, Edge-to-Node (E2N) layer, Node-to-Graph (N2G) layer, and a final fully connected (Linear) layer. The number of the E2E layers can be any number greater than or equal to zero. On the other hand, there is one E2N layer and one N2G layer. The number of convolution filters and number of nodes in these layers are among the many hyperparameters that have to be tuned.

The BrainNetCNN takes in any connectivity matrix directly as an input and outputs behavioral or demographic predictions. Kawahara et al. (2017) used this model for predicting age and neurodevelopmental outcomes from structural connectivity data. BrainNetCNN consists of four types of layers: Edge-to-Edge (E2E) layer, Edge-to-Node (E2N) layer, Node-to-Graph (N2G) layer and a final fully connected (linear) layer. The first three types of layers are specially designed layers introduced in the BrainNetCNN. The final fully connected layer is the same as that used in FNNs.

The Edge-to-Edge (E2E) layer is a convolution layer using cross-shaped filters (Figure 2). The cross-shaped filter is applied to each element of the input matrix. Thus, for each filter, the E2E layer takes in a *N* × *N* matrix and outputs a *N* × *N* matrix. The number of E2E layer is arbitrary and is a tunable hyperparameter. The outputs of the final E2E layer are inputs to the E2N layer. The E2N layer is similar to the E2E layer, except that the cross-shaped filter is applied to only the diagonal entries of the input matrix. Thus, for each filter, the E2N layer takes in a *N* × *N* matrix and outputs a *N* × 1 vector. There is one E2N layer for BrainNetCNN. The outputs of the E2N layer are the inputs to the Node-to-Graph (N2G) layer. The N2G layer is simply a fully connected hidden layer similar to the a FNN’s hidden layer. Finally, the outputs of the N2G layer are linearly summed by the final fully connected layer to provide a final set of prediction values.

### 2.7 Graph convolutional neural network (GCNN)

Standard convolution applies to data that lies on a Euclidean grid (e.g., images). Graph convolution exploits the graph Laplacian in order to generalize the concept of standard convolution to data lying on nodes connected together into a graph. This allows the extension of the standard CNN to graph convolutional neural networks (GCNNs; Defferrard et al., 2016; Bronstein et al., 2017; Kipf and Welling, 2017). There are many different ways that GCNN can be applied to neuroimaging data (Kipf and Welling, 2017; Ktena et al., 2018; Zhang et al., 2018). Here we considered the innovative GCNN developed by Kipf and Welling (2017) and extended to neuroimaging data by Parisot and colleagues (Parisot et al., 2017, 2018). Figure 3 illustrates this approach.

**Figure 3.**
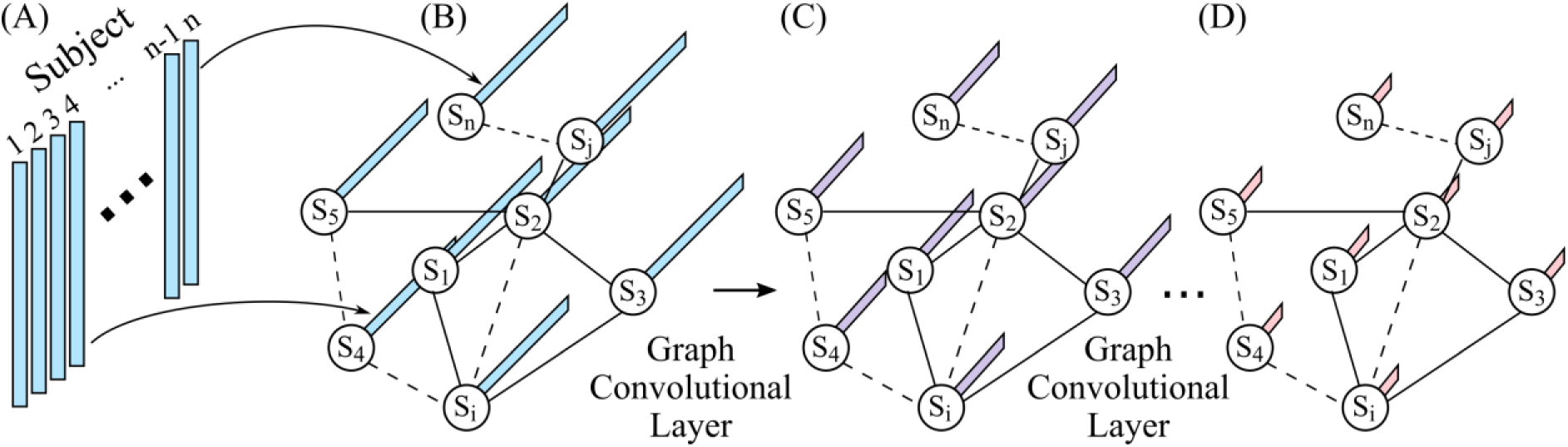
Schematic of a graph convolution neural network. (GCNN; Parisot et al., 2017, 2018). This particular GCNN takes in vectorized RSFC matrices of all subjects as input and outputs behavioral (or demographic) prediction of all subjects. (A) Vectorized FC of all subjects (subject 1 to subject n). (B) The input of GCNN is a graph, where each node represents a subject and is associated with the vectorized FC from the corresponding subject in (A). An edge in the graph represents the similarity between two subjects. Here, the similarity is defined in terms of the similarity of the subjects’ RSFC matrices. (C) Output of the first graph convolutional layer. Graph convolutional layer extends standard convolution to graph convolution. Each node is associated with a vector, whose length is determined by the number of filters in the first graph convolutional layer. (D) Final output of GCNN after one or more graph convolutional layers. Each node contains the predicted behavioral measure(s).

The input to an FNN (Figure 1) or a BrainNetCNN (Figure 2) is the RSFC data of a single subject. By contrast, the GCNN takes in data (e.g., vectorized RSFC) of *all* subjects as input and outputs behavioral (or demographic) predictions of *all* subjects (Parisot et al., 2017, 2018). In other words, data from the training, validation, and testing sets are all input into the GCNN at the same time. To avoid leakage of information across training, validation and test sets, masking of data is applied during the calculation of the loss function and gradient descent.

More importantly, the graph in GCNN does not represent connectivity matrices (like in BrainNetCNN). Instead, each node represents a subject and edges are determined by the similarity between subjects. This similarity is problem dependent. For example, in the case of autism spectrum disorder (ASD) classification, similarity between two subjects is defined based on sex, sites and RSFC, i.e., two subjects are more similar if they have the same sex, from the same site and have similar RSFC patterns (Parisot et al., 2017, 2018). The use of sex and sites in the graph definition were particularly important for this specific application, since ASD is characterized by strong sex-specific effects and the database included data from multiple unharmonized sites (Di Martino et al., 2014).

Similar to the original studies (Parisot et al., 2017, 2018), we utilized vectorized RSFC (lower triangular entries of the RSFC matrix) of all subjects as inputs to the GCNN. Edges between subjects were defined based on Pearson’s correlation between lower triangular portions of RSFC matrices.

### 2.8 HCP training, validation and testing

For the HCP dataset, 20-fold cross-validation was performed. The 953 subjects were divided into 20 folds, such that family members were not split across folds. Inner-loop cross-validation was performed for hyperparameter tuning. More specifically, for a given test fold, cross-validation was performed on the remaining 19 folds with different hyperparameters. The best hyperparameters were then used to train on the 19 folds. The trained model was then applied to the test fold. This was repeated for all 20 test folds.

In the case of kernel regression, there was only one single hyperparameter λ (that controls the *l*_2_ regularization; see Appendix A.1). A separate hyperparameter was tuned for each fold and each behavioral measure separately based on a grid search over the hyperparameter.

In the case of the DNNs, there was a large number of hyperparameters, e.g., number of layers, number of nodes, number of training epochs, dropout rate, optimizer (e.g., stochastic gradient or ADAM), weight initialization, activation functions, regularization, etc. GCNN also has additional hyperparameters tuned, e.g., definition of the graph and graph Laplacian estimation. Therefore, instead of training a separate DNN for each behavioral measure, a single FNN (or BrainNetCNN or GCNN) was trained for all 58 behavioral measures. The reason is that tuning hyperparameters separately for each behavioral measure would be too time consuming. We note that the joint prediction of multiple behavioral measures might not be a disadvantage for the DNNs and might even potentially improve prediction performance because of shared structure among target behavioral variables (Rahim et al., 2017). Furthermore, we tried to tune each DNN (FNN, BrainNetCNN or GCNN) for only fluid intelligence, but the performance for fluid intelligence prediction was not better than predicting all 58 behavioral measures simultaneously.

Furthermore, a proper inner-loop 20-fold cross-validation would involve tuning the hyperparameters for each DNN 20 times (once for each split of the data into training-test folds), which was computationally prohibitive. Thus, for each DNN (FNN, BrainNetCNN and GCNN), we tuned the hyperparameters once, using the first split of the data into training-test folds, and simply re-used the optimal hyperparameters for the remaining training-test splits of the data. Such a procedure biases the prediction performance in favor of the DNNs (relative to kernel regression), so the results should be interpreted accordingly (see Discussion). Such a bias is avoided in the UK Biobank dataset (see below). Further details about DNN hyperparameters are found in Appendix A2.

As is common in the FC-based prediction literature (Finn et al., 2015), model performance was evaluated based on the Pearson’s correlation between predicted and actual behavioral measures across subjects within each test fold. Furthermore, since certain behavioral measures were correlated with motion (Siegel et al., 2017), age, sex, and motion (FD) were regressed from the behavioral measures from the training and test folds (Kong et al., 2019; Li et al., 2019). Regression coefficients were estimated from the training folds and applied to the test folds. Mean absolute error (MAE) and coefficient of determination (COD) will also be reported.

### 2.9 UK Biobank training, validation and testing

The large UK Biobank dataset allowed us the luxury of splitting the 8868 subjects into training (N = 6868), validation (N = 1000) and test (N = 1000) sets, instead of employing an inner-loop cross-validation procedure like in the HCP dataset. Care was taken so that the distributions of various attributes (sex, age, fluid intelligence and pairs matching) were similar across training, validation and test sets.

Hyperparameters were tuned using the training and validation sets. The test set was only utilized to evaluate the final prediction performance. A separate DNN was trained for each of the four behavioral and demographic measures. Thus, the hyperparameters were tuned independently for each behavioral/demographic measure. Further details about DNN hyperparameters are found in Appendix A2. Initial experiments using a single neural network to predict all four measures simultaneously (like in the HCP dataset) did not appear to improve performance and so was not further pursued. In the case of kernel regression, the hyperparameter λ was tuned using the validation set based on a grid search over the hyperparameter.

Like before, prediction accuracies for age, fluid intelligence and pairs matching were evaluated based on the Pearson’s correlation between predicted and actual measures across subjects within the test set. Since the age prediction literature often used mean absolute error (MAE) as an evaluation metric (Liem et al., 2017; Cole et al., 2018; Varikuti et al., 2018), we included MAE as an evaluation metric. For completeness, we also computed MAE for pairs matching and fluid intelligence.

In the case of sex, accuracy was defined as the fraction of participants whose sex was correctly predicted. Like before, we regressed age, sex and motion from fluid intelligence and pairs matching measures in the training set and apply the regression coefficients to the validation and test sets. When predicting age and sex, no regression was performed. Coefficient of determination (COD) for age, pairs matching and fluid intelligence will also be reported in the Supplemental Material.

### 2.10 Deep neural network implementation

The DNNs were implemented using Keras (Chollet, 2015) or PyTorch (Paszke et al., 2017) and run on NVIDIA Titan Xp GPU using CUDA. Our implementation of BrainNetCNN and GCNN were based on GitHub code from the original papers (Kawahara et al., 2017; Kipf and Welling, 2017). Our implementations achieved similar results as the original implementations when using the toy datasets and hyperparameters provided by the original GitHub implementations. More details about hyperparameter tuning can be found in Appendix A2.

### 2.11 Statistical tests

For the HCP dataset, we performed 20-fold cross-validation, yielding a prediction accuracy for each test fold. To compare two algorithms, the corrected resampled t-test was performed (Nadeau and Bengio, 2003; Bouckaert and Frank, 2004). The corrected resampled t-test corrects for the fact that the accuracies across test folds were not independent. In the case of the UK Biobank, there was only a single test fold, so the corrected resampled t-test could not be applied. Instead, when comparing correlations from two algorithms, the Steiger’s Z-test was utilized (Steiger, 1980). When comparing MAE, a two-tailed paired sample t-test was performed. When comparing prediction accuracies for sex, the McNemar’s test was utilized (McNemar, 1947).

### 2.12 Scaling of prediction performance as a function of sample size

The large UK Biobank dataset allowed us to explore the effect of sample size on predicting fluid intelligence. The test set (N = 1000) was the same as before to allow for meaningful comparisons. We considered 100, 500, 1000, 2000, 3000, 4000, 5000 and 6000 and 7868 subjects for training and validation. The case of 7868 subjects was identical to the analysis from the previous sections.

In the case of 3000, 4000, 5000 and 6000 subjects, the validation set comprised the same set of 1000 subjects as in the previous sections. The training set was obtained by randomly sampling the appropriate number of subjects from the original training set of 6868 participants. For example, in the case of 3000 training and validation subjects, we randomly sampled 2000 training subjects from the original training set. However, the training subjects were selected so that the distribution of fluid intelligence matched the distributions of the validation and test sets.

In the case of 100, 500, 1000 and 2000 subjects, we split the participants with a 3:1 ratio. For example, in the case of 100 subjects, there were 75 training and 25 validation subjects. Like before, the participants were randomly selected but we ensured the distributions of fluid intelligence in the training and validation sets were similar to the distribution of the test set.

The hyperparameter tuning for the three DNNs and kernel regression was the same as in previous sections. See Appendices A1 and A2 for more details.

### 2.13 Control analysis

We repeated our analyses using hyperparameters as close as possible to the original BrainNetCNN hyperparameters (provided by the BrainNetCNN code repository; Kawahara et al., 2017) and original GCNN hyperparameters (provided by the GCNN code repository; Parisot et al., 2017; 2018). In the case of FNN, we utilized hyperparameters as close as possible to the FC90net baseline in the BrainNetCNN paper (Kawahara et al., 2017).

### 2.14 Data and code availability

This study utilized publicly available data from the HCP (https://www.humanconnectome.org/) and UK Biobank (https://www.ukbiobank.ac.uk/). The 400 cortical ROIs (Schaefer et al., 2018) can be found here (https://github.com/ThomasYeoLab/CBIG/tree/master/stable_projects/brain_parcellation/Schaefer2018_LocalGlobal). The kernel regression and DNNs code utilized in this study can be found here (https://github.com/ThomasYeoLab/CBIG/tree/master/stable_projects/predict_phenotypes/He2019_KRDNN). The trained models for the UK Biobank dataset can also be found in the above GitHub link. The code was reviewed by one of the co-authors (MN) before merging into the GitHub repository to reduce the chance of coding errors.

## 3. Results

### 3.1 HCP behavioral prediction

Figure 4 shows the prediction accuracy (Pearson’s correlation coefficient) averaged across 58 HCP behavioral measures and 20 test folds. Statistical tests were performed between kernel regression and the three DNNs (see Methods). False discovery rate (q < 0.05) was applied to correct for multiple comparisons correction.

**Figure 4.**
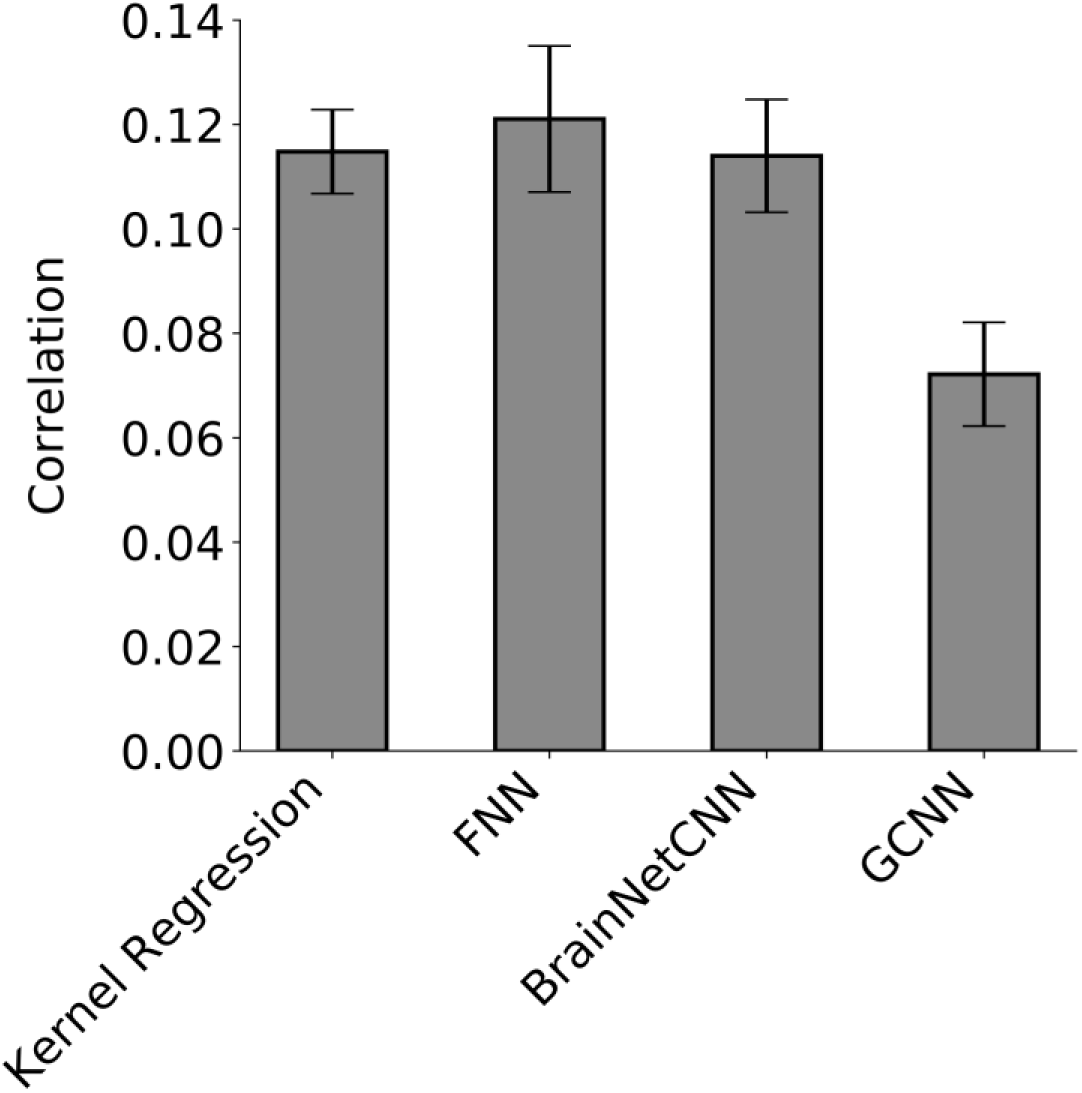
Prediction accuracy (Pearson’s correlation coefficient) averaged across 58 HCP behavioral measures and 20 test folds. Correlation was computed for each test fold and each behavior, and then averaged across the 58 behaviors. Bars show mean across test folds. Error bars show standard error of model performance across cross-validation folds. Kernel regression and FNN performed the best. There was no statistical difference between kernel regression and FNN or BrainNetCNN. Kernel regression was statistically better than GCNN (p = 3e-4).

FNN achieved the highest average prediction accuracy with Pearson’s correlation r = 0.121 ± 0. 063 (mean ± std). On the other hand, kernel regression achieved an average prediction accuracy of r = 0.115 ± 0. 036 (mean ± std). However, there was no statistical difference between FNN and kernel regression (p = 0.60). Interestingly, BrainNetCNN (r = 0.114 ± 0. 048) and GCNN (r = 0.072 ± 0. 044) did not outperform FNN, even though the two DNNs were designed for neuroimaging data. KRR was significantly better than GCNN (p = 3e-4), but not BrainNetCNN (p = 0.93).

For completeness, Figures 5, S2, and S3 show the behavioral prediction accuracies for all 58 behavioral measures. Figures S4 to S7 show the scatterplots of predicted versus actual values for 13 cognitive measures. Kernel regression was significantly better than FNN for predicting grip strength (p = 2.65e-4) and significantly better than GCNN for predicting picture matching vocabulary (p = 6.91e-5). No other difference survived the FDR correction.

**Figure 5.**
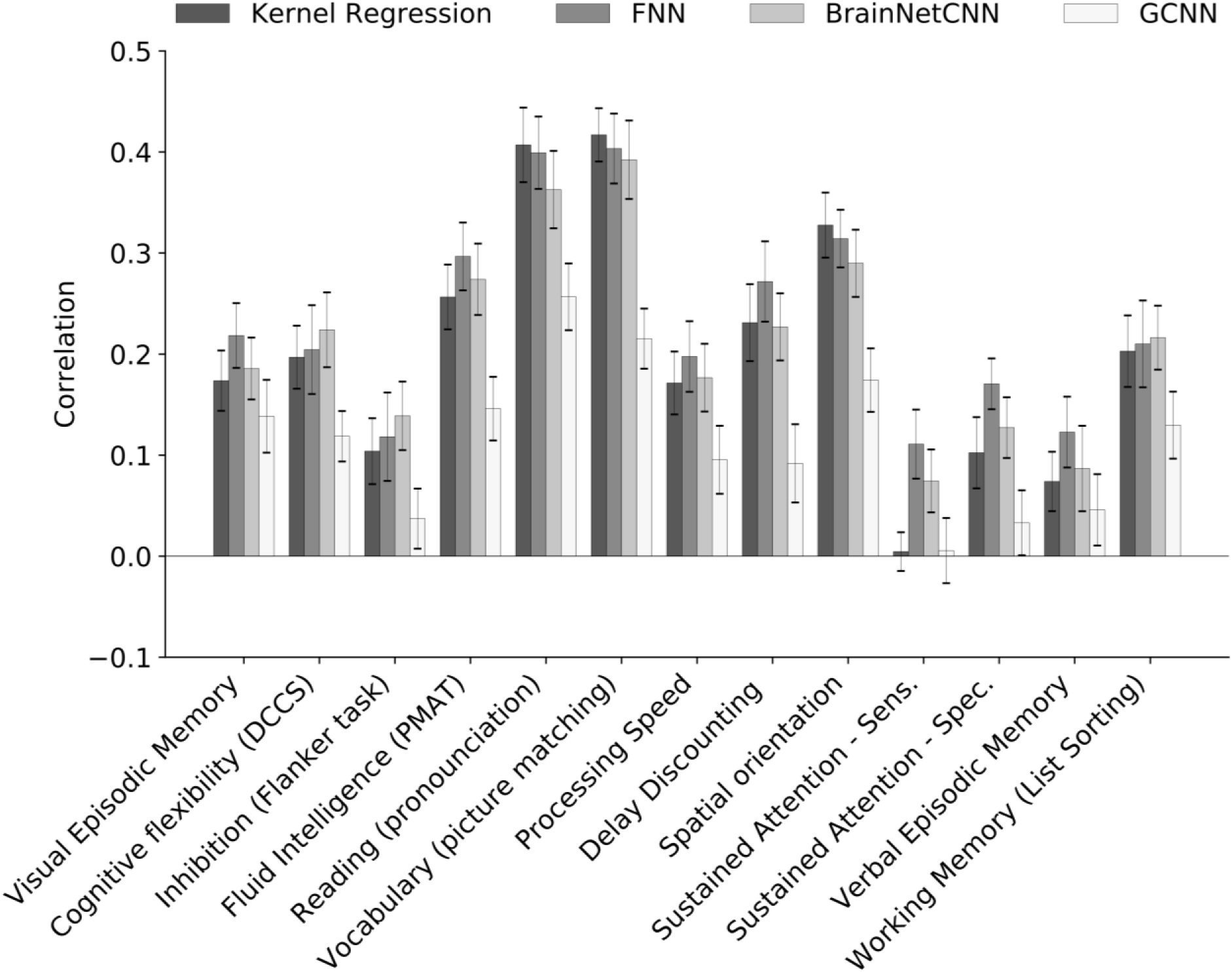
Prediction accuracies (Pearson’s correlation coefficient) in a curated set of 13 HCP cognitive measures averaged across 20 test folds. Correlation was computed for each test fold and each behavior. Bars show mean across test folds. Error bars show standard errors of model performance across cross-validation folds. Prediction accuracies of the remaining 45 behavioral measures are found in Figures S2 and S3.

Similar conclusions were obtained when using mean absolute error (Figure 6) and coefficient of determination (Figure S8) as measures of prediction performance.

**Figure 6.**
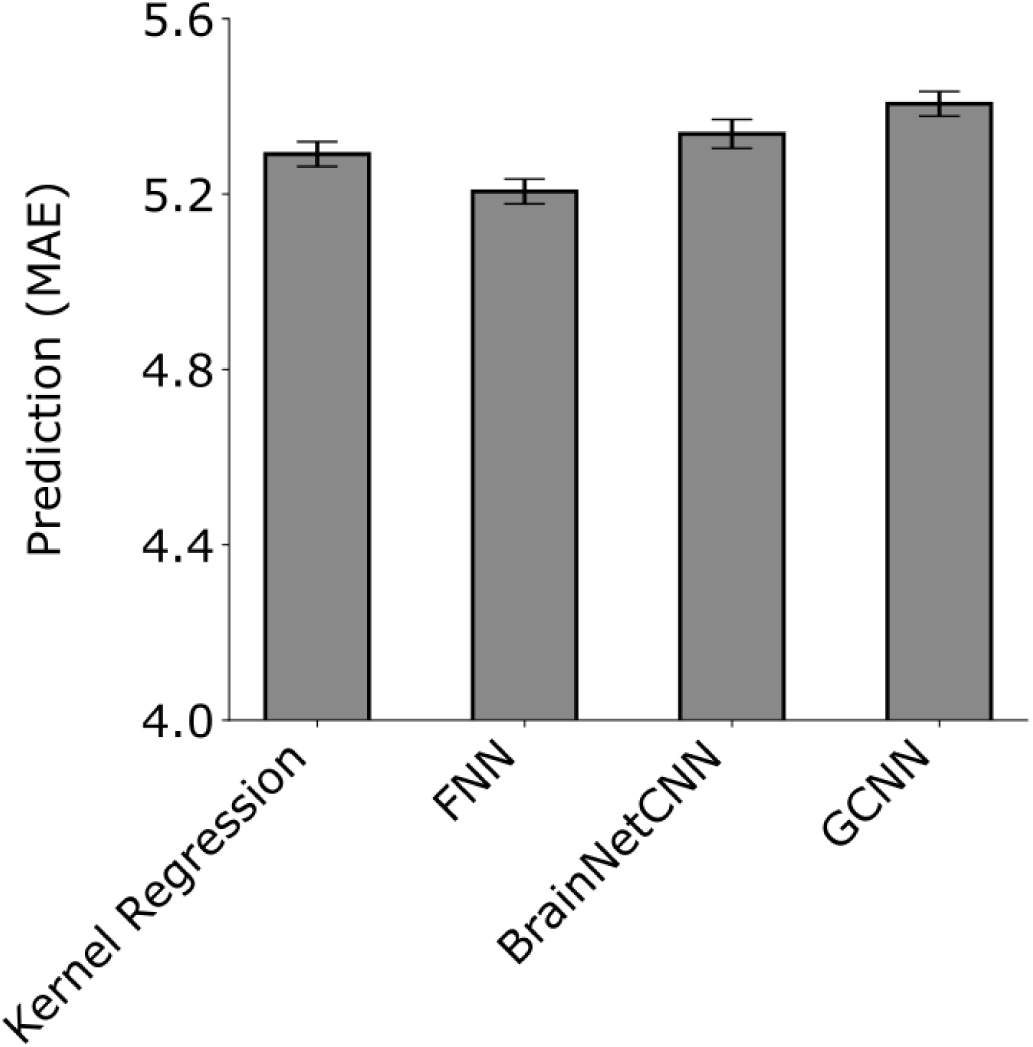
Prediction MAE averaged across 58 HCP behavioral measures and 20 test folds. Lower is better. MAE was computed for each test fold and each behavior and then averaged across the 58 behaviors. Bars show the mean across test folds. Error bars show standard error of model performance across cross-validation folds. There was no statistical difference between kernel regression and all DNNs after correcting for multiple comparisons.

### 3.2 UK Biobank behavioral and demographics prediction

Table 1 and Figure 7 show the prediction performance of sex, age, pairs matching and fluid intelligence. Figure S9 shows the scatterplots of predicted versus actual values for age, pairs matching and fluid intelligence. Kernel regression, FNN, and GCNN achieved the highest accuracy for sex prediction. Kernel regression performed the best for fluid intelligence and age (measured using Pearson’s correlation). BrainNetCNN performed the best for age (measured using MAE) and pairs matching.

**Figure 7.**
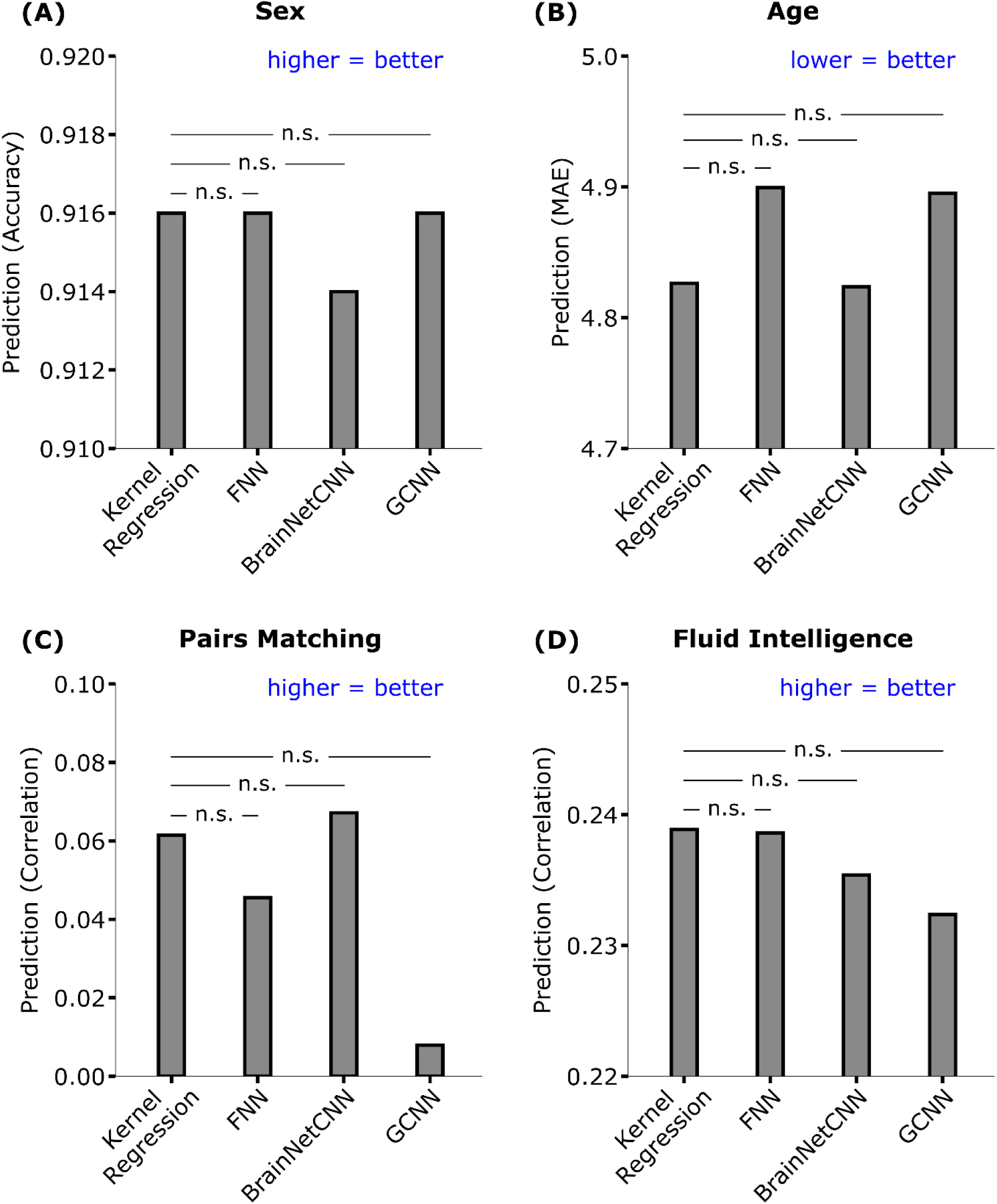
Prediction performance of four behavioral and demographic measures in the UK Biobank. For age (MAE), lower values imply better performance. For all the other measures, larger values imply better performance. The horizontal lines represent statistical tests between kernel regression and the DNNs. “n.s” stands for not significant after FDR (q < 0.05) correction.

Statistical tests were performed between kernel regression and the three DNNs (see Methods). False discovery rate (q < 0.05) was applied to correct for multiple comparisons correction. There was no statistical difference between kernel regression and the DNNs for all behavioral and demographic measures.

Interestingly, the GCNN achieved poor performance in the case of pairs matching (Pearson’s correlation r = 0.008), although it was not statistically worse than kernel regression. Upon further investigation, we found that GCNN achieved an accuracy of r = 0.106 in the UK Biobank validation set. When using the initial set of hyperparameters (before hyperparameter tuning using HORD), GCNN achieved accuracies of r = 0.047 and r = 0.056 in the validation and test sets respectively. Overall, this suggests that the hyperparameter tuning overfitted the validation set, despite the rather large sample size.

Similar conclusions were obtained when using mean absolute error (MAE) as a performance measure for fluid intelligence and pairs matching (Table 2 and Figure S10), or when using coefficient of determination (COD) as a performance measure for age, pairs matching and fluid intelligence (Table S2).

**Table 2.** Prediction MAE of pairs matching and fluid intelligence in the UK Biobank. Lower values imply better performance. **Bold** indicates the best performance. We note that simply predicting the median of the pairs matching value in the training set would have yielded an MAE of 0.400, which was better than kernel regression and all DNNs.

| **Model** | **Pairs matching** | **Fluid intelligence** |
| --- | --- | --- |
| Kernel regression | 0.551 | **1.608** |
| FNN | 0.567 | 1.613 |
| BrainNetCNN | 0.553 | 1.610 |
| GCNN | **0.497** | 1.612 |
| Median | 0.400 | 1.656 |

### 3.3 Effect of sample size on predicting fluid intelligence in the UK Biobank

Figure 8 shows the prediction performance (Pearson’s correlation) of fluid intelligence in the UK Biobank as the training and validation sample sizes were varied, while the same test set of 1000 subjects was used throughout. All algorithms performed poorly with 100 subjects but improved with more subjects. There was more than 300% improvement when increasing the sample size from 100 to 1000 subjects and more than 35% improvement when increasing the sample size from 1000 to 5000 subjects. However, the improvement tapered off from 5000 to 7868 subjects. GCNN was highly volatile as the sample size was varied, suggesting its sensitivity to particular choices of training and validation subjects. Kernel regression was competitive across all sample sizes. Similar conclusions were obtained when MAE was used as a performance metric (Figure S11).

**Figure 8.**
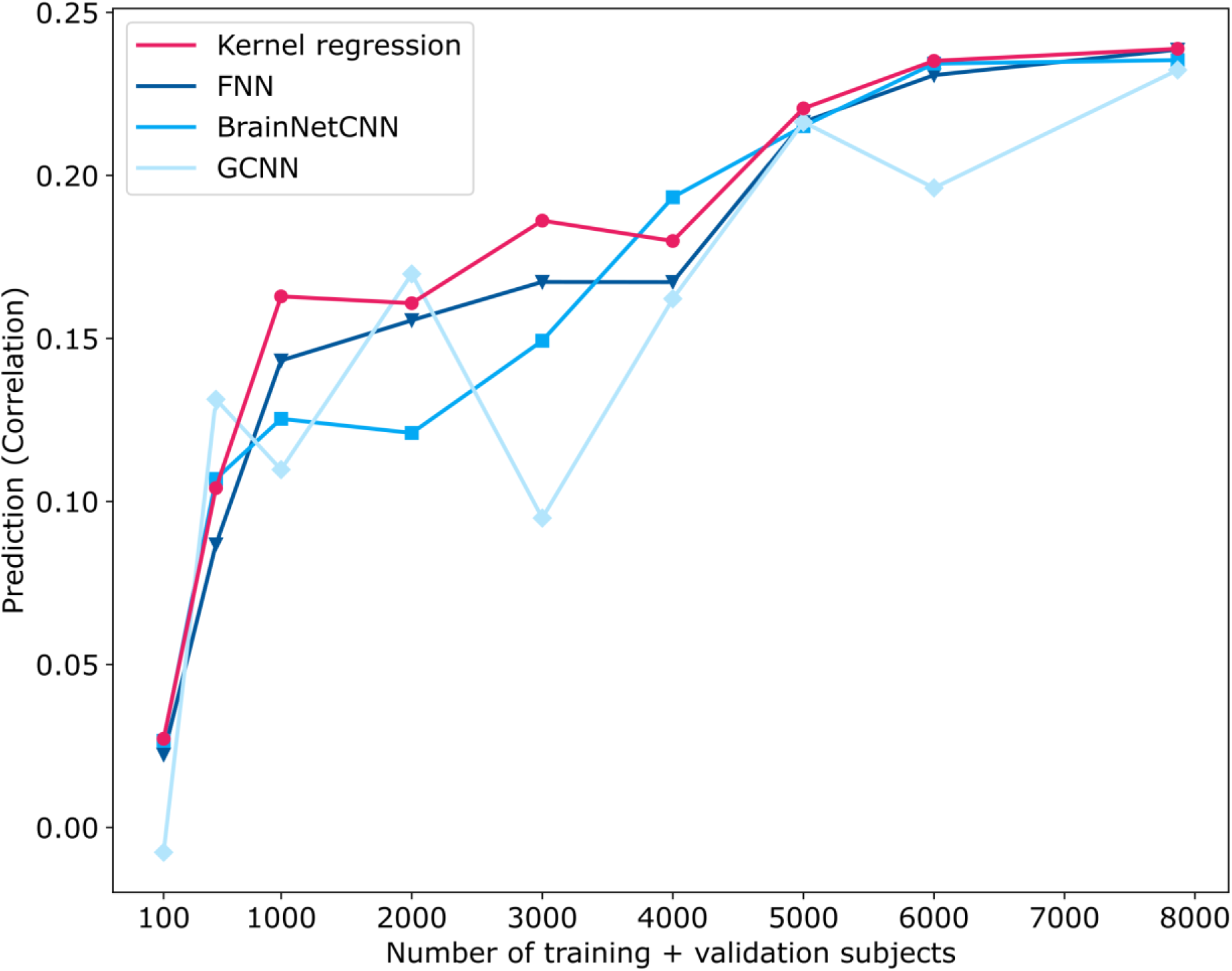
Prediction performance (Pearson’s correlation coefficient) of fluid intelligence in the UK Biobank dataset with different number of training and validation subjects. The performance of all algorithms generally increased with more training and validation subjects. In the case of 100, 500, 1000 and 2000 subjects, 3/4 of the subjects were used for training and 1/4 of the subjects were used for validation. In the remaining cases, 1000 subjects were used for validation, while the remaining subjects were used for training. For all cases, test set comprised the same set of 1000 subjects. Kernel regression was highly competitive across all sample sizes. See Figure S11 for MAE results.

### 3.4 Control analysis

Tables S3 and S4 show the performance of the DNNs using hyperparameters from the original publications (Kawahara et al., 2017; Parisot et al., 2017; 2018) versus our tuned hyperparameters. The performance of our hyperparameters compared favorably to the performance of the original hyperparameters. This is not surprising, since our hyperparameters were obtained by tuning using the datasets shown in this paper.

**Table 3.** FNN architecture and hyperparameters for HCP and UK Biobank. Under “Model structure”, the numbers represent the number of nodes at each fully connected layer. For example, “256, 96, 256, 58” represents a 4-layer FNN with 256, 96, 256 and 58 nodes.

| Dataset | Predicting | Model architecture | Optimizer |
| --- | --- | --- | --- |
| HCP | 58 behaviors | 223, 128, 192, 58 | SGD |
| UK Biobank | Sex | 3, 2 | SGD |
|  | Age | 9, 1 | SGD |
|  | Pairs matching | 415, 437, 1 | SGD |
|  | Fluid intelligence | 318, 357, 1 | SGD |

### 3.5 Computational costs

Kernel regression has a close-form solution (Appendix A1) and only one hyperparameter, so the computational cost is extremely low. For example, kernel regression training and grid search of 32 hyperparameter values in the UK Biobank validation set took about 20 minutes (single CPU core) for one behavioral measure. This is one reason why we considered kernel regression instead of other slower classical approaches (e.g., support vector regression or elastic net) requiring iterative optimization. On the other hand, FNN training and tuning of hyperparameters in the UK Biobank validation set took around 80 hours (single GPU) for one behavioral measure, excluding the manhours necessary for the manual tuning.

## 4. Discussion

In this study, we showed that kernel regression and DNNs achieved similar performance in RSFC-based prediction of a wide range of behavioral and demographic measures across two large-scale datasets totaling almost 10,000 participants. Furthermore, FNN performed as well as the two DNNs that were specifically designed for connectome data. Given comparable performance between kernel regression and the DNNs and the significantly greater computational costs associated with DNNs, our results suggest that kernel regression might be more suitable than DNNs in some neuroimaging applications.

### 4.1 Potential reasons why DNNs did not outperform kernel regression for RSFC-based prediction

There are a few potential reasons why DNNs did not outperform kernel regression in our experiments on RSFC-based behavioral prediction. First, while the human brain is nonlinear and hierarchically organized (Deco et al., 2011; Breakspear, 2017; Wang et al., 2019), such a structure might not be reflected in the RSFC matrix in a way that was exploitable by the DNNs we considered. This could be due to the measurements themselves (Pearson’s correlations of rs-fMRI timeseries), the particular representation (N x N connectivity matrices) or particular choices of DNNs, although we again note that BrainNetCNN and GCNN were specifically developed for connectome data.

Second, given the much larger datasets used in computer vision and natural language processing (Chelba et al., 2014; Russakovsky et al., 2015), it is possible that there was not enough neuroimaging data (even in the UK Biobank) to fully exploit DNNs. However, our experiments show that kernel regression was highly competitive across all sample sizes from 100 to 7898 subjects. In fact, all approaches (except GCNN) improved at almost lockstep with greater sample size, suggesting that even larger sample sizes might equally benefit both DNNs and kernel regression.

Third, it is well-known that hyper-parameter settings and architectural details can impact the performance of DNNs. Thus, it is possible that the benchmark DNNs we implemented in this work can be further optimized. However, we do not believe this would alter our conclusions for two reasons. First, for some measures (e.g., sex classification in the UK Biobank), we were achieving performance at or near the state-of-the-art. Second, an earlier version of this paper relied completely on manual tuning of hyperparameters. In the current version of this paper, we utilized an automatic algorithm to tune a subset of hyperparameters for the UK Biobank experiments (Appendix A2), yielding essentially the same conclusions.

It is also worth pointing out that while deep learning has won several predictive modeling challenges, these have mostly involved image segmentation (Choi et al., 2016, Kamnitsas et al., 2017a, Hongwei Li et al., 2018). The success of DNNs has been less clear in other neuroimaging challenges. For example, in the 2019 ABCD challenge to predict fluid intelligence from structural MRI, kernel regression was the winner, beating other deep learning algorithms (Mihalik et al., 2019). Similarly, in the recent TADPOLE challenge to predict Alzheimer’s Disease progression (Marinescu et al., 2018), the top entry did not utilize deep learning (https://tadpole.grand-challenge.org/Results/).

### 4.2 Hyperparameters

There are significantly more hyperparameters in DNNs compared with classical machine learning approaches. For example, for a fixed kernel (e.g., correlation metric in our study), kernel regression has one single regularization parameter. Even with a nonlinear kernel (e.g. radial basis function), there would only be two hyperparameters. This is in contrast to DNNs, where there are easily more than ten hyperparameters.

Because of the large number of hyperparameters, most applications involving DNNs currently require some level of manual hyperparameter tuning. Therefore, we suggest that manual hyper-parameter tuning should be performed within a training-validation-test framework (like in our UK Biobank experiments), rather than a nested (inner-loop) cross-validation framework (like in HCP experiments). The reason is that within a nested (inner-loop) cross-validation framework, information from tuning one fold might leak to another fold (via the person tuning the hyperparameters).

To elaborate, recall that we divided the HCP dataset into 20 folds. We tuned the hyperparameters of the DNNs using folds 2 to 20 and applied the trained DNNs to fold 1. Since fold 1 was not used in tuning the hyperparameters, the performance of the DNNs in fold 1 was unbiased. However, when fold 2 became the test fold, we utilized the same hyperparameters to train using folds 1, 3 to 20. This is problematic because fold 2 was originally utilized to tune the hyperparameters, so consequently the performance of the DNNs in test fold 2 was inflated.

One could try to independently tune the hyperparameters for each fold independently. However, complete independence between folds is unlikely because the person performing the manual tuning cannot possibly forget his/her tuning experience with the other folds. As such, this will yield overly optimistic results.

On the other hand, the test set in the UK Biobank was only utilized after the hyperparameters have been determined from the training and validation sets. Therefore, the performance of the DNNs was unbiased. It is worth noting that our motivation for advocating the training-validation-test framework is to prevent overly optimistic results in the test set, but does not necessarily eliminate overfitting. For example, in the case of pairs matching in the UK Biobank, our tuning procedure overfitted on the validation set, yielding poor performance in the test set (Table 1). Thus, overfitting was “caught” in the test set, which highlights the benefits of adopting a training-validation-test framework.

Finally, we note that there are generally too many DNN hyperparameters (and design choices) to be listed in a paper. In fact, there were hyperparameters too complex to completely specify in this paper. However, we have made our code publicly available, so researchers can refer to the code for the exact hyperparameters. We encourage future neuroimaging DNN studies to also make their code publicly available.

### 4.3 Prediction performance in the literature

Comparing our prediction performance with the literature is difficult because of different datasets, sample sizes, cross-validation procedures and de-confounding strategies. For example, we regressed age, sex, and motion (FD) from the behavioral measures, but other studies might not perform any regression or use a different set of regressors. Nevertheless, we believe that our prediction performance is generally consistent with the literature.

As mentioned earlier, our sex prediction accuracy of 91.6% in the UK Biobank is among the best in the literature. For example, Ktena and colleagues (2018) reported a sex prediction accuracy of around 80% when using 55 x 55 functional connectivity matrices from 2500 UK Biobank subjects. On the other hand, Chekroud and colleagues (2016) reported sex prediction accuracy of 93% when using cortical thickness and subcortical volumes of 1566 subjects from the Brain Genomics Superstruct Project (Holmes et al., 2015).

In the case of fluid intelligence, our prediction accuracies (Pearson’s correlation) ranged from around 0.257 to 0.297 (excluding GCNN which performed poorly) in the HCP dataset. Although earlier RSFC-based behavioral prediction studies have reported high fluid intelligence prediction accuracy in the HCP dataset (Finn et al., 2015), newer studies using more subjects reported lower accuracies comparable with our results. For example, Dubois and colleagues (2018) reported a prediction accuracy (Pearson’s correlation) of 0.27 for fluid intelligence in the HCP dataset. On the other hand, Greene and colleagues (2018) reported a prediction accuracy (Pearson’s correlation) of 0.17 for fluid intelligence in the HCP dataset (but only using data from a single resting-fMRI session). Thus, our prediction accuracies for fluid intelligence is consistent with the literature.

In the case of age prediction, we achieved a prediction accuracy (Pearson’s correlation) of 0.6 and an MAE of 4.8 in the UK Biobank dataset. Comparing these results with the literature is difficult because of sensitivity to age range in the dataset. For example, many studies utilized either lifespan (Cole et al., 2017; Liem et al., 2017) or developmental (Sturmfels et al., 2018; Nielsen et al., 2019) cohorts, while the UK Biobank comprised older adults (more than 45 years old). Furthermore, many studies preferred to use structural MRI, instead of RSFC, for predicting age (Cole et al., 2017; Sturmfels et al., 2018; Varikuti et al., 2018). Liem and colleagues (2017) achieved MAEs ranging from 5.25 to 5.99 when using RSFC for predicting age in a lifespan dataset comprising 2354 subjects, which was worse than our MAE. On the other hand, their prediction accuracies (Pearson’s correlation) ranged from 0.79 to 0.93, which was better than our prediction accuracy (Pearson’s correlation). Overall, this suggests that our prediction performance is probably comparable with other RSFC studies, although we emphasize that comparing age prediction performance across datasets is non-trivial.

It is important to mention that prediction performance was poor for a number of target variables across all four prediction algorithms. For example, in the case of pairs matching in the UK Biobank dataset, predicting the median of the training set yielded lower MAE than all four models, suggesting that pairs matching is not an easily predictable trait using RSFC. Therefore, it might not be meaningful to compare the models for pairs matching. On the other hand, we note that for both age and fluid intelligence prediction, all four models performed better than predicting the median of the training set. Similarly, sex prediction was a lot better than chance, given that there were roughly equal number of males and females in the dataset. For these three target variables (age, sex and fluid intelligence), all four models exhibited very similar performance.

It is also worth noting that the poor average COD in the HCP dataset is consistent with the literature. For example, of the 58 behavioral measures, 48 of them were also utilized in the HCP MegaTrawl (https://db.humanconnectome.org/megatrawl/). For the 300-dimensional group-ICA results, HCP MegaTrawl achieved an average COD of -0.177 (original data space), while kernel regression in the current study achieved an average COD of -0.0875. Overall, this suggests that certain target variables are not easily predicted using RSFC.

### 4.4 Limitations and caveats

Although the current study suggests that kernel regression and DNNs achieved similar performance for RSFC-based behavioral prediction, it is possible that other DNNs (we have not considered) might outperform kernel regression. Furthermore, our study focused on the use of N x N static RSFC matrices for behavioral prediction. Other RSFC features, such as dynamic RSFC features (Calhoun et al., 2014; Preti et al., 2017; Liégeois et al., 2019), in combination with DNNs might potentially yield better performance (Hongming Li et al., 2018; Khosla et al., 2019).

We also note that our evaluation procedure was performed on the HCP and UK Biobank datasets independently. Therefore, we expect the reported prediction performance to be maintained if new participants were recruited in the respective studies under the same experimental conditions (e.g., no scanner upgrade, same population, same acquisition protocol and preprocessing, etc). However, the reported prediction performance would likely drop if the trained models (from the UK Biobank or HCP) were applied to other datasets (Arbabshirani et al., 2017; Woo et al., 2017). At this point, it is unclear which approach (kernel regression, FNN, BrainNetCNN or GCNN) would generalize better to a completely new dataset. This is obviously an active area of research given the increasing number of large-scale publicly available brain imaging datasets.

## 5. Conclusion

By using a combined sample of nearly 10,000 participants, we showed that kernel regression and three types of DNN architectures achieved similar performance for RSFC-based prediction of a wide range of behavioral and demographic measures. Overall, our study suggests that kernel regression might be just as effective as DNNs for certain neuroimaging applications, while incurring significantly less computational costs.

## Acknowledgment

We like to thank the anonymous reviewers for the very helpful feedback. We would also like to thank Christine Annette, Taimoor Akhtar, Li Zhenhua for their help on the HORD algorithm. This work was supported by Singapore MOE Tier 2 (MOE2014-T2-2-016), NUS Strategic Research (DPRT/944/09/14), NUS SOM Aspiration Fund (R185000271720), Singapore NMRC (CBRG/0088/2015), NUS YIA and the Singapore National Research Foundation (NRF) Fellowship (Class of 2017). Our research also utilized resources provided by the Center for Functional Neuroimaging Technologies, P41EB015896 and instruments supported by 1S10RR023401, 1S10RR019307, and 1S10RR023043 from the Athinoula A. Martinos Center for Biomedical Imaging at the Massachusetts General Hospital. Our computational work was partially performed on resources of the National Supercomputing Centre, Singapore (https://www.nscc.sg). The Titan Xp GPUs used for this research were donated by the NVIDIA Corporation. This research has been conducted using the UK Biobank resource under application 25163 and Human Connectome Project, WU-Minn Consortium (Principal Investigators: David Van Essen and Kamil Ugurbil; 1U54MH091657) funded by the 16 NIH Institutes and Centers that support the NIH Blueprint for Neuroscience Research; and by the McDonnell Center for Systems Neuroscience at Washington University.

## Appendix

### A1. Kernel Regression

In this section, we describe kernel regression in detail (Liu et al., 2007; Murphy, 2012). The kernel matrix *K* encodes the similarity between pairs of subjects. Motivated by Finn and colleagues (2015), the *i*-th row and *j*-th column of the kernel matrix is defined as the Pearson’s correlation between the *i*-th subject’s vectorized RSFC and *j*-th subject’s vectorized RSFC (considering only the lower triangular portions of the RSFC matrices). The behavioral measure *y_i_* of subject *i* can be written as:

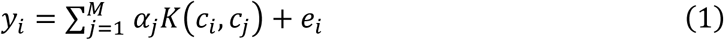

where *c_i_* is the vectorized RSFC of the *i*-th subject, *K*(*c_i_*, *c_j_*) is the element at *i*-th row and *j*-th column of kernel matrix, *M* is the total number of training subjects, *e_i_* is the noise term and α_j_ is the trainable weight. The goal of kernel regression is to find an optimal set of α. To achieve this goal, we maximize the penalized likelihood function:

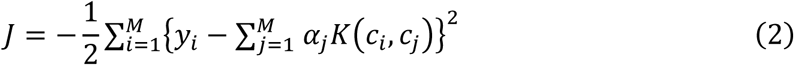

with respect to **α** = [α_1_, α_2_, …, α*_M_*]*^T^*. To avoid overfitting, a *l*_2_ regularization (i.e., kernel ridge regression) can be added, so the resulting optimization problem becomes:

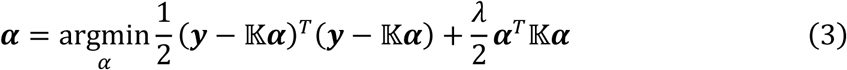

where 𝕂 is the *M* × *M* kernel matrix, **y** = [*y*_1_, *y*_2_, …, *y_M_*]*^T^* and λ is a hyperparameter that controls the *l*_2_ regularization. By solving equation (3) with respect to *α*, we can predict a test subject’s behavioral measure *y_s_* as:

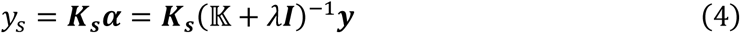

where ***K_s_*** = [*K*(*c_s_*, *c*_1_), *K*(*c_s_*, *c*_2_),…, *K*(*c_s_*, *c_M_*)].

In the case of the HCP, λ was selected via inner-loop cross-validation. In the case of the UK biobank, λ was tuned on the validation set. For sex prediction in the UK Biobank, for each continuous prediction *y_sex_*, the participant was classified as male or female based on whether it was larger or smaller than the threshold. We tuned the threshold to obtain the best accuracy in the UK Biobank validation dataset and used this threshold in the test set.

### A2 More details of deep neural networks

In this section, we describe further details of our DNN implementation.

- For GCNN, we adopted Keras code from the GCNN GitHub repository (https://github.com/tkipf/keras-gcn; Kipf and Welling, 2017). We made some minor modifications to the code, e.g., the modified code directly loaded the graph adjacency matrix, instead of loading the edges and generating the adjacency matrix. As another example, our graph convolution layer loaded the graph matrix as parameters rather than as an input. However, we emphasized that the core functionalities (e.g., graph convolution) remained unchanged. As a sanity check, we applied our modified code to the original toy data using the original hyperparameters provided by the original GitHub repository. Our results were comparable to the original implementation (Table S5).
- The original BrainNetCNN implementation used the Caffe framework (https://github.com/jeremykawahara/ann4brains; Kawahara et al., 2017). We re-implemented BrainNetCNN in Keras and PyTorch following the original Caffe code as closely as possible. The Keras version was applied to the HCP data, while the PyTorch version was applied to the UK Biobank data. The reason for this inconsistency was that after our experiments with the HCP dataset using Keras, we realized that the Keras framework yielded slightly different results each time the code was run. This was apparently a well-known issue of the framework. As such, we decided to implement a second version in PyTorch, which was then applied to the UK Biobank. As a sanity check, we applied both implementations (Keras and PyTorch) to the original toy data using the original hyperparameters provided by the original GitHub repository. Our implementations achieved comparable results with the original implementation (Table S6).
- In the case of the FNN, since this is just a generic feedforward neural network, so we implemented using default libraries in Keras and PyTorch. The Keras version was applied to the HCP data, while the PyTorch version was applied to the UK Biobank data. The reason for this inconsistency is the same as the previous bullet point.
- Representative learning curves for the HCP dataset are shown in Figure S12. Learning curves for the UK Biobank are shown in Figures S13 to S15. The training curves showed good accuracy/error, suggesting that we are not underfitting to the data. The validation curves were plateauing, suggesting that we were not stopping too early in our training. Since the validation and test curves were progressing in almost lockstep (except for certain instances of GCNN), our stopping criterion (based on the peaks of the validation curves) was reasonable. For most behavioral measures, there were relatively big gaps between the training and validation/test curves, suggesting overfitting. However, we have already deployed several standard strategies to reduce overfitting, including dropout, L2-regularization/weight-decay and batch-normalization.

In the case of the HCP dataset:

- For all three DNNs, all behavioral measures were z-normalized based on training data. The loss function was mean squared error (MSE). Optimizer was stochastic gradient descent (SGD). With the MSE loss, the output layer has 58 nodes (FNN and BrainNetCNN) or filters (GCNN).
- In the case of the main results (Figures 4, 5, S2 and S3), the hyperparameters were tuned manually by trial-and-error. Since each test fold was of size 47 or 48, we simply set 48 to be the batch size (except GCNN, which utilized the whole dataset in a single mini-batch). We initialized with a default set of hyperparameters (e.g., learning rate=0.01, dropout rate=0.5, number of filter/nodes=32) and then tuned the optimizer (learning rate, momentum, and learning rate decay), layer structure (number of layers, number of nodes/filters), dropout rate, regularization and weight initialization. There was no fixed order for hyperparameter tuning. We generally started by tuning the layer structure, followed by the optimizer and then other hyperparameters. For GCNN, we also tuned the graph-related hyperparameters at the beginning of the tuning process.
- Final FNN structure is shown in Table 3. Dropout of 0.6 was added before each fully-connected layer. L2 regularization of 0.02 was added for layer 2.
- Final BrainNetCNN structure is shown in Table 4. Dropout of 0.4 was added after E2N layer. LeakyReLU (Maas et al., 2013) with alpha of 0.3 was used as the activation function for the first three layers.
- Final GCNN structure is shown in Table 5. Dropout of 0.3 was added for each layer. L2 regularization of 8e-4 was added for layer 1. The nodes of the graph corresponded to subjects. Edges were constructed based on Pearson’s correlation between subjects’ vectorized RSFC. The graph was thresholded by only retaining edges with top 5% correlation (across the entire graph). However, this might result in a disconnected graph. Therefore, the top five correlated edges of each node were also retained (even if these edges were not among the top 5% correlated edges). The graph convolution filters were estimated using a 5-degree Chebyshev polynomial (Defferrard et al., 2016).

In the case of the UK Biobank:

**Table 4.** BrainNetCNN architecture and hyperparameters for HCP and UK Biobank. Under “Model structure”, the numbers represent the number of filters or nodes at each layer. For example, “15, 93, 106, 2” represents a BrainNetCNN with 15 filters for the E2E layer, 93 filters for the E2N layer, 106 filters (nodes) for the N2G layer and 2 nodes in the final fully connected layer. All BrainNetCNNs follow the same layer order: E2E, E2N, N2G and then a final fully connected layer.

| Dataset | Predicting | Model architecture | Optimizer |
| --- | --- | --- | --- |
| HCP | 58 behaviors | 18, 19, 84, 58 | SGD |
| UK Biobank | Sex | 38, 58, 7, 2 | SGD |
|  | Age | 22, 79, 91, 1 | SGD |
|  | Pairs matching | 27, 29, 54, 1 | SGD |
|  | Fluid intelligence | 40, 60, 41, 1 | SGD |

**Table 5.** GCNN architecture and hyperparameters for HCP and UK Biobank. Under “Model structure”, the numbers represent the number of filters for each graph convolutional layer. For example, “64, 1” represents a 2-layer GCNN with 64 and 1 filters respectively.

| Dataset | Predicting | Model architecture | Optimizer |
| --- | --- | --- | --- |
| HCP | 58 behaviors | 256, 58 | SGD |
| UK Biobank | Sex | 71, 2 | Adam |
|  | Age | 10, 1 | SGD |
|  | Pairs matching | 3, 1 | Adam |
|  | Fluid intelligence | 72, 1 | Adam |

- For all three DNNs, model ensemble was used to improve final test result: for each DNN and each behavior, five models were trained separately (with different random initializations). The predictions were averaged across the five models yielding a final prediction. All four behavioral measures were z-normalized based on training data. The loss function for sex prediction was cross entropy, i.e., the output layer for sex prediction have 2 nodes (FNN and BrainNetCNN) or filters (GCNN). The loss function was MSE for the other three measures. The output layer for these three measures have 1 node (FNN and BrainNetCNN) or filter (GCNN). Adam (Kingma and Ba, 2015) or SGD were used. See details in Tables 3, 4 and 5.
- For all three DNNs, we utilized the HORD algorithm (Regis et al., 2013; Ilievski et al., 2017, Eriksson et al., 2019) to assist in hyperparameter tuning using the UK Biobank validation dataset. For each DNN, the HORD algorithm automatically tuned the DNN hyperparameters within user-specified ranges of various hyperparameters. Not all hyperparameters were tuned by HORD because the speed and performance of HORD worsened when too many hyperparameters were tuned. Therefore, we determined several hyperparameters based on our previous manual tuning experience, i.e. momentum = 0.9, batch size = 128 (except GCNN’s batch size is 1 as it loads all data at once), weight initialization = Xavier uniform (PyTorch) or Glorot uniform (Keras), Chebyshev polynomial basis filters with degree of 1 for GCNN.
- For FNN, we tuned the number of layers (2 to 4 layers), number of nodes for each layer (2 to 512 nodes), dropout rate (0 to 0.8), starting learning rate (1e-2 to 1e-4), weight decay rate (1e-3 to 1e-7), and epochs to decrease learning rate (10 to 200 epochs) using HORD.
- For BrainNetCNN, we tuned the number of filters for e2e (2 to 48 filters), e2n (2 to 96 filters), and n2g layers (2 to 128 nodes), dropout rate (0 to 0.8), learning rate (1e-2 to 1e-4), weight decay rate (1e-3 to 1e-7), and epochs to decrease the learning rate (10 to 200 epochs) using HORD.
- For GCNN, we tuned the number of filters for GCNN layer (2 to 128 filters), methods to generate graph adjacency matrix, dropout rate (0 to 0.8), L2 regularization rate (1e-3 to 1e-7), and learning rate (1e-2 to 1e-4) using HORD.
- For all DNNs, model was tuned for each behavior separately. Tables 3, 4 and 5 show the final DNN structures and hyperparameters.
- Final FNN structure is shown in Table 3. For FNN, dropout of 0.00275/0.309/0.285/0.526 (for sex/age/pairs matching/fluid intelligence respectively) were added before each fully-connected layer. L2 regularization of 0.02 was added for layer 2. Weight decay of 2.662e-4/2.799e-5/1.141e-6/1.425e-4 (for sex/age/pairs matching/fluid intelligence respectively) were applied to the weights of all fully connected layers.
- Final BrainNetCNN structure is shown in Table 4. For BrainNetCNN, dropout of 0.463/0.573/0.264/0.776 (for sex/age/pairs matching/fluid intelligence respectively) were added after the E2E, E2N, and N2G layers. LeakyReLU was replaced by linear activation for all four models.
- Final GCNN structure is shown in Table 5. Dropout of 0.0150/0.316/0.308/0.555 (for sex/age/pairs matching/fluid intelligence respectively) were added before the first and second hidden layers. L2 regularization of 3.344e-4/9.181e-7/4.716e-7/7.183e-4 (for sex/age/pairs matching/fluid intelligence respectively) were added for layer 1. The nodes of the graph corresponded to subjects. Edges were constructed based on Pearson’s correlation between subjects’ vectorized RSFC. Thresholding of the graph was tuned separately for each behavior or demographic measure. For pairs matching prediction, the top five correlated edges of each node were retained. For age, sex and fluid intelligence prediction, the graph was thresholded by only retaining edges with top 5% correlation (across the entire graph). Furthermore, the top five correlated edges of each node were also retained (even if these edges were not among the top 5% correlated edges). The graph convolution filters for all four GCNNs were estimated by a 1-degree Chebyshev polynomial (Defferrard et al., 2016).

---

**Figure S1.**
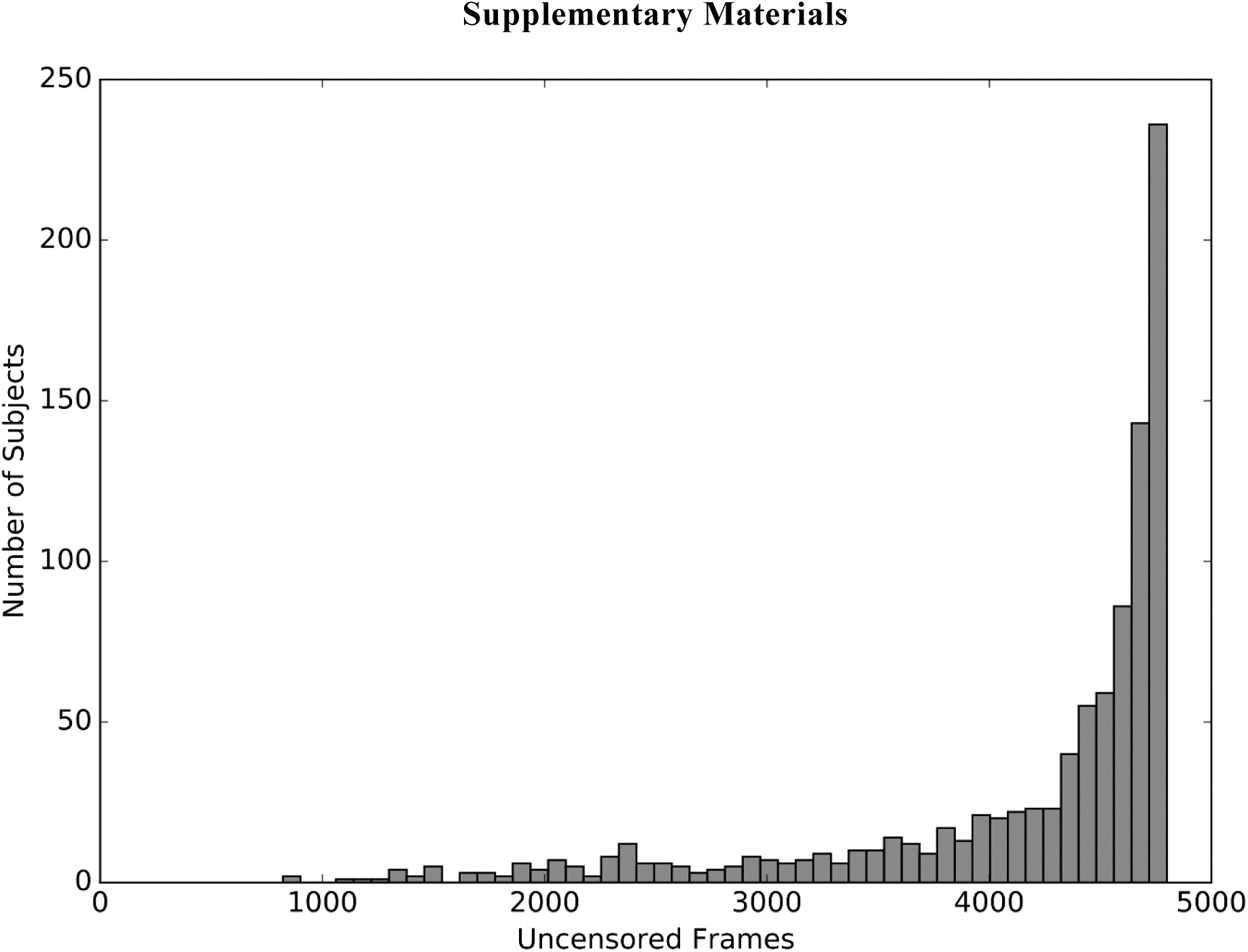
Distribution of the number of uncensored frames across 953 HCP subjects. The subject with the least uncensored frames had 822 frames, which corresponded to almost ten minutes of data.

**Table S1.**
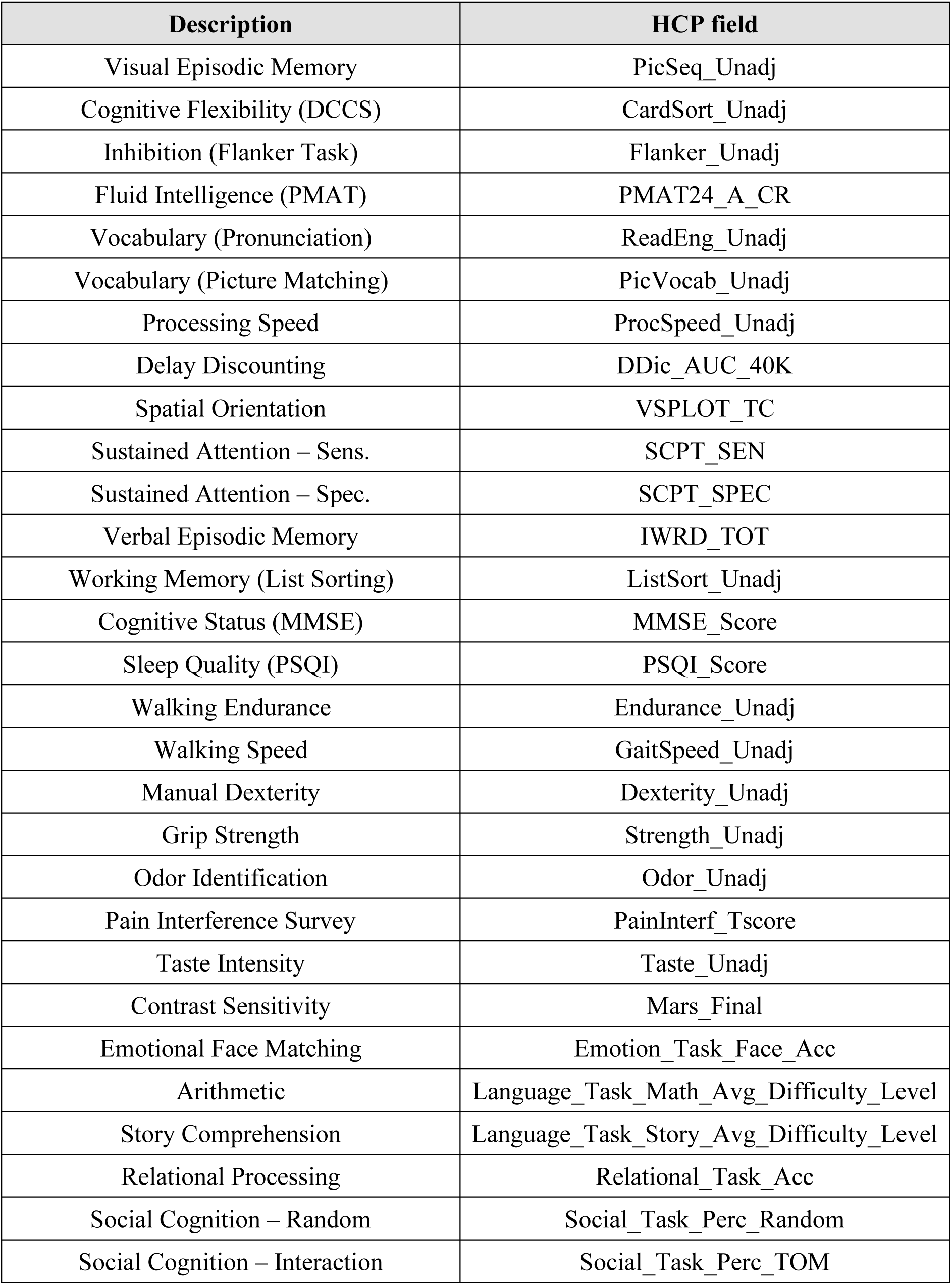
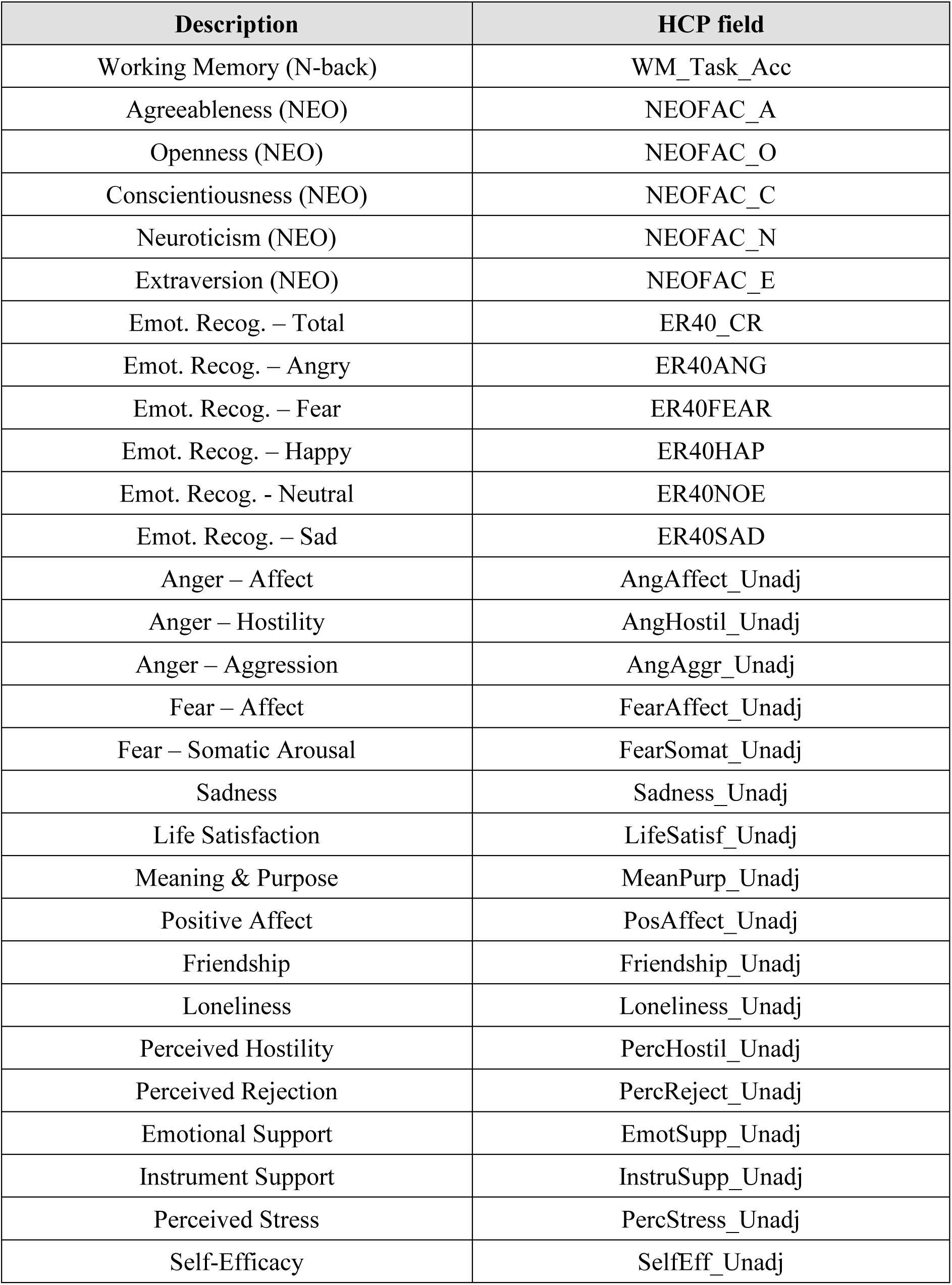
Table showing original HCP variable names and corresponding descriptive labels used in the manuscript.

**Figure S2.**
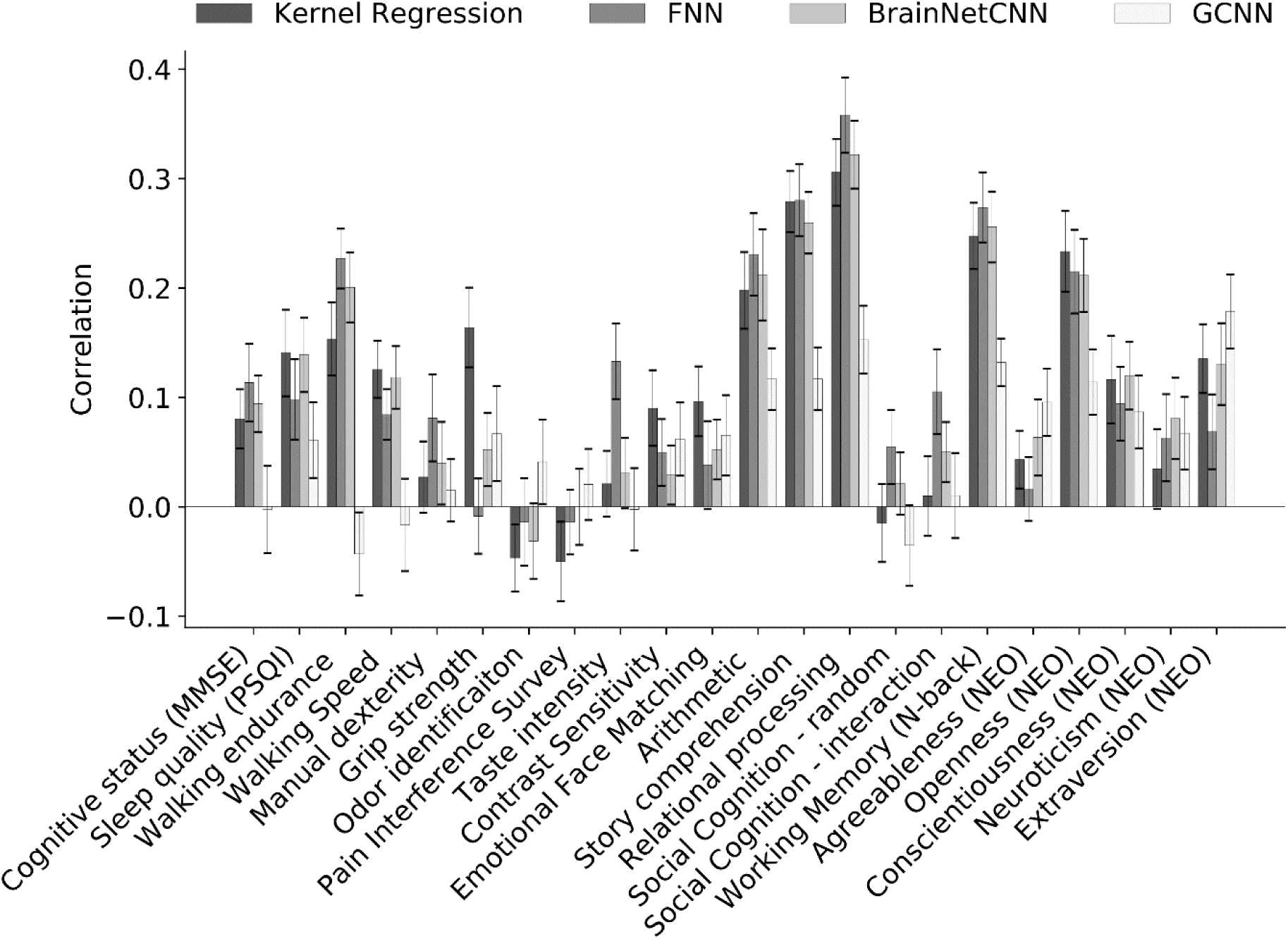
Prediction accuracy (Pearson’s correlation coefficient) of 22 HCP measures averaged across 20 test folds. Correlation was computed for each test fold and each behavior. Bars show mean across test folds. Error bars show standard errors.

**Figure S3.**
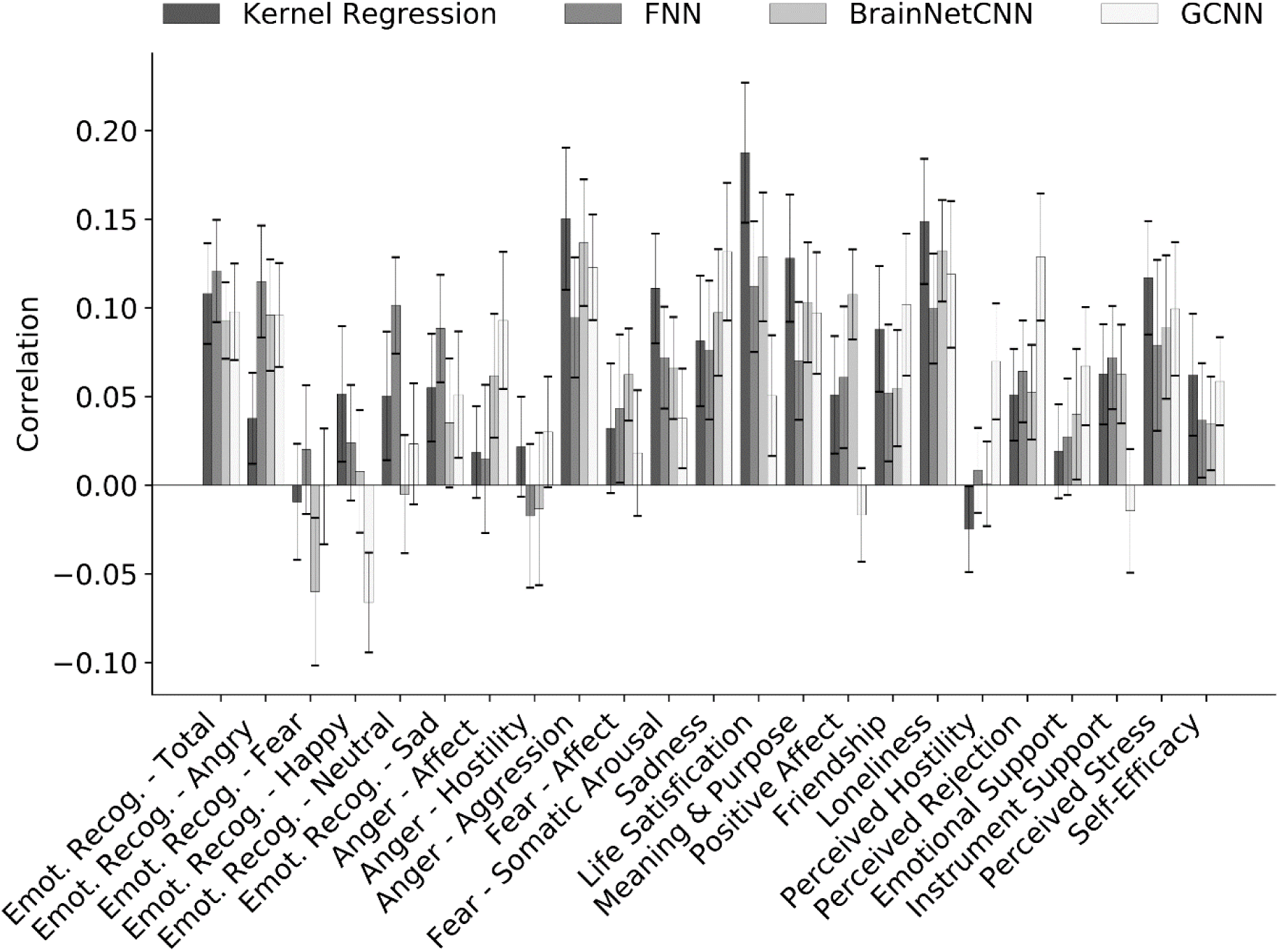
Prediction accuracy (Pearson’s correlation coefficient) of 23 HCP cognitive measures averaged across 20 test folds. Correlation was computed for each test fold and each behavior. Bars show mean across test folds. Error bars show standard errors.

**Figure S4.**
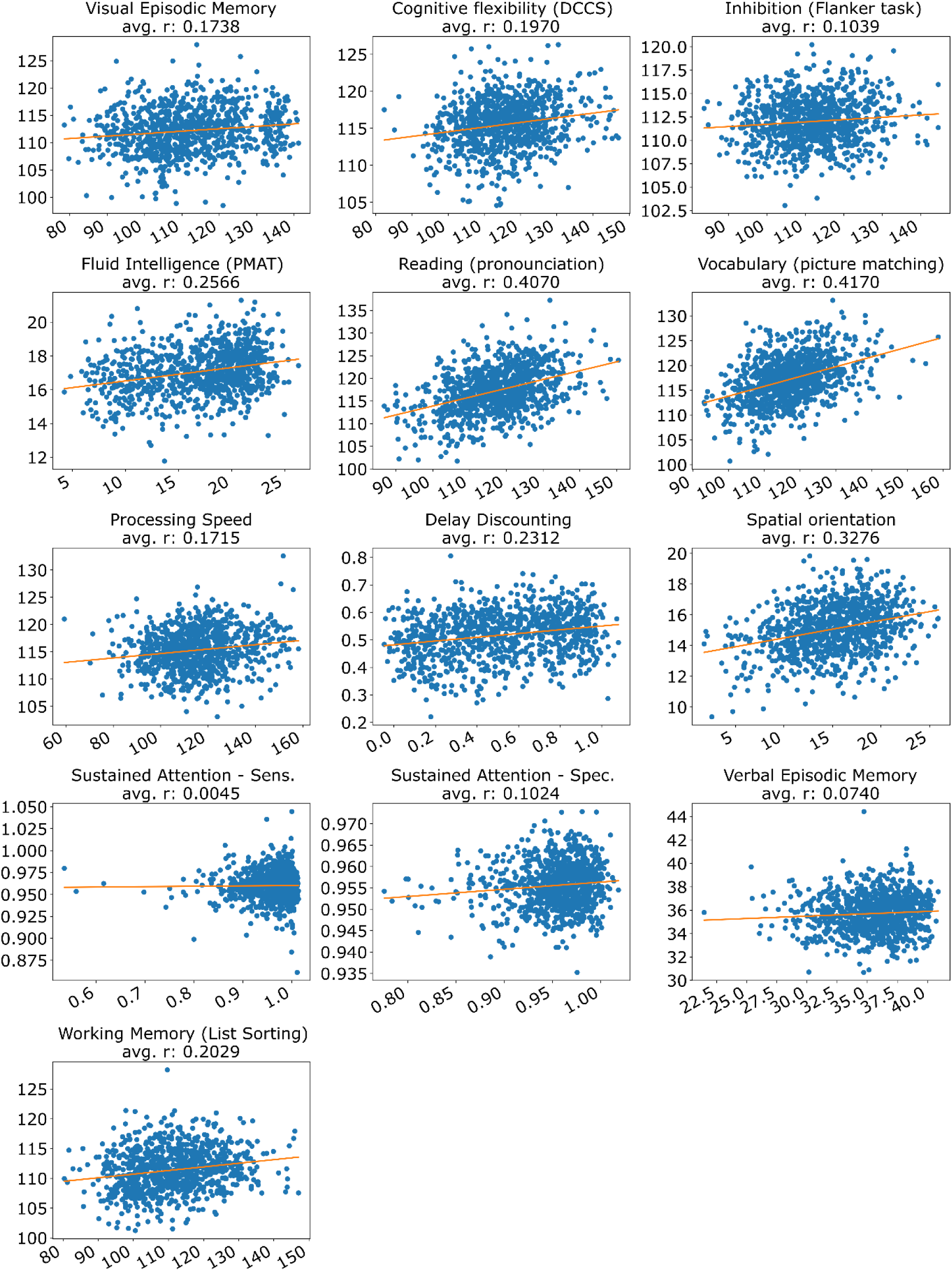
Scatterplots of predicted (kernel regression) versus actual values for a curated set of 13 HCP cognitive measures for all 953 subjects. X-axis is the actual value. Y-axis is the predicted value. Each dot represents one subject. The orange line shows the best fit line obtained by least squares regression. The “avg. r” in each subplot title is the average Pearson’s correlation coefficient across 20 test folds, which was also reported in Figure 5.

**Figure S5.**
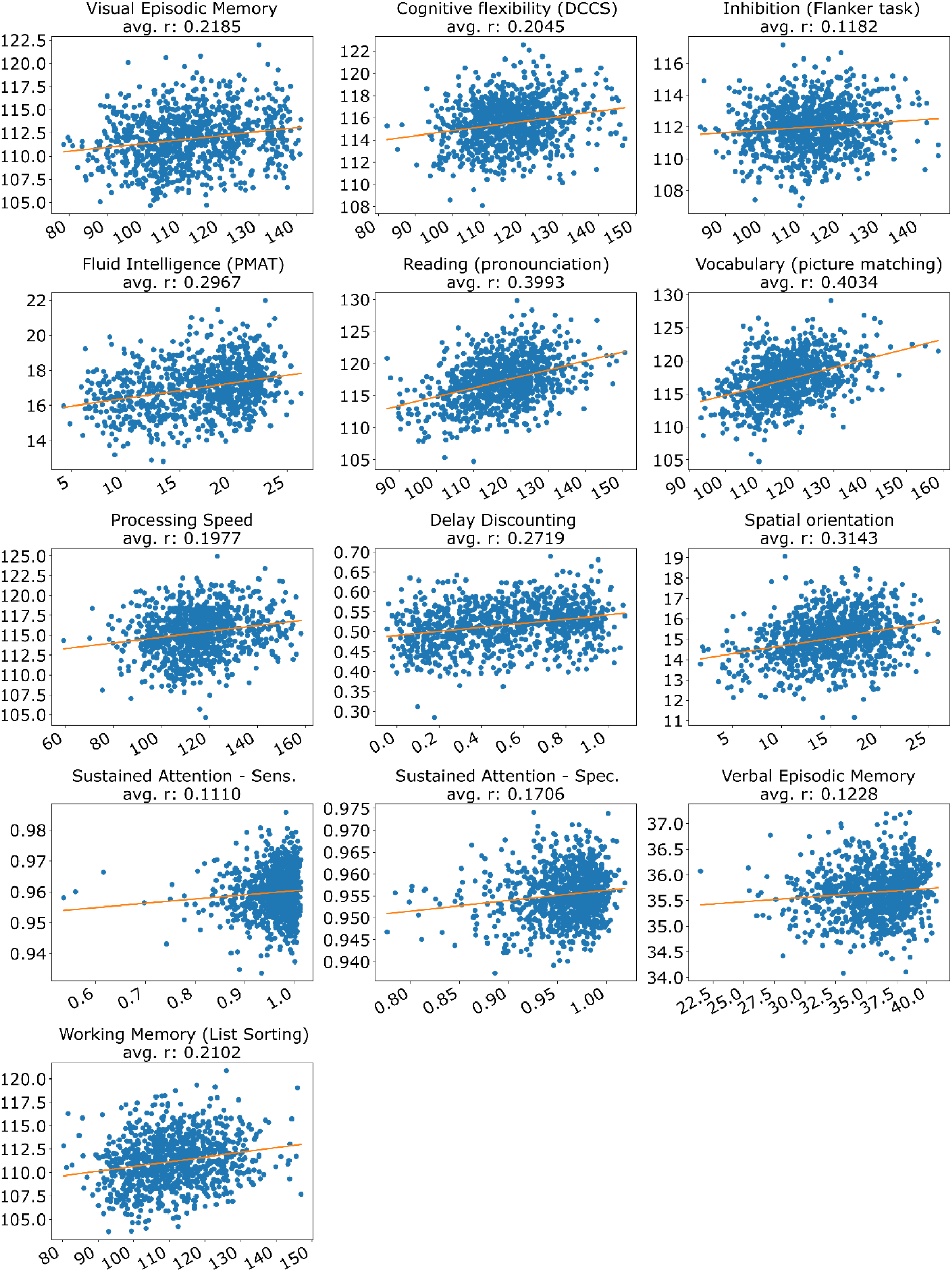
Scatterplots of predicted (FNN) versus actual values for a curated set of 13 HCP cognitive measures for all 953 subjects. X-axis is the actual value. Y-axis is the predicted value. Each dot represents one subject. The orange line shows the best fit line obtained by least squares regression. The “avg. r” in each subplot title is the average Pearson’s correlation coefficient across 20 test folds, which was also reported in Figure 5.

**Figure S6.**
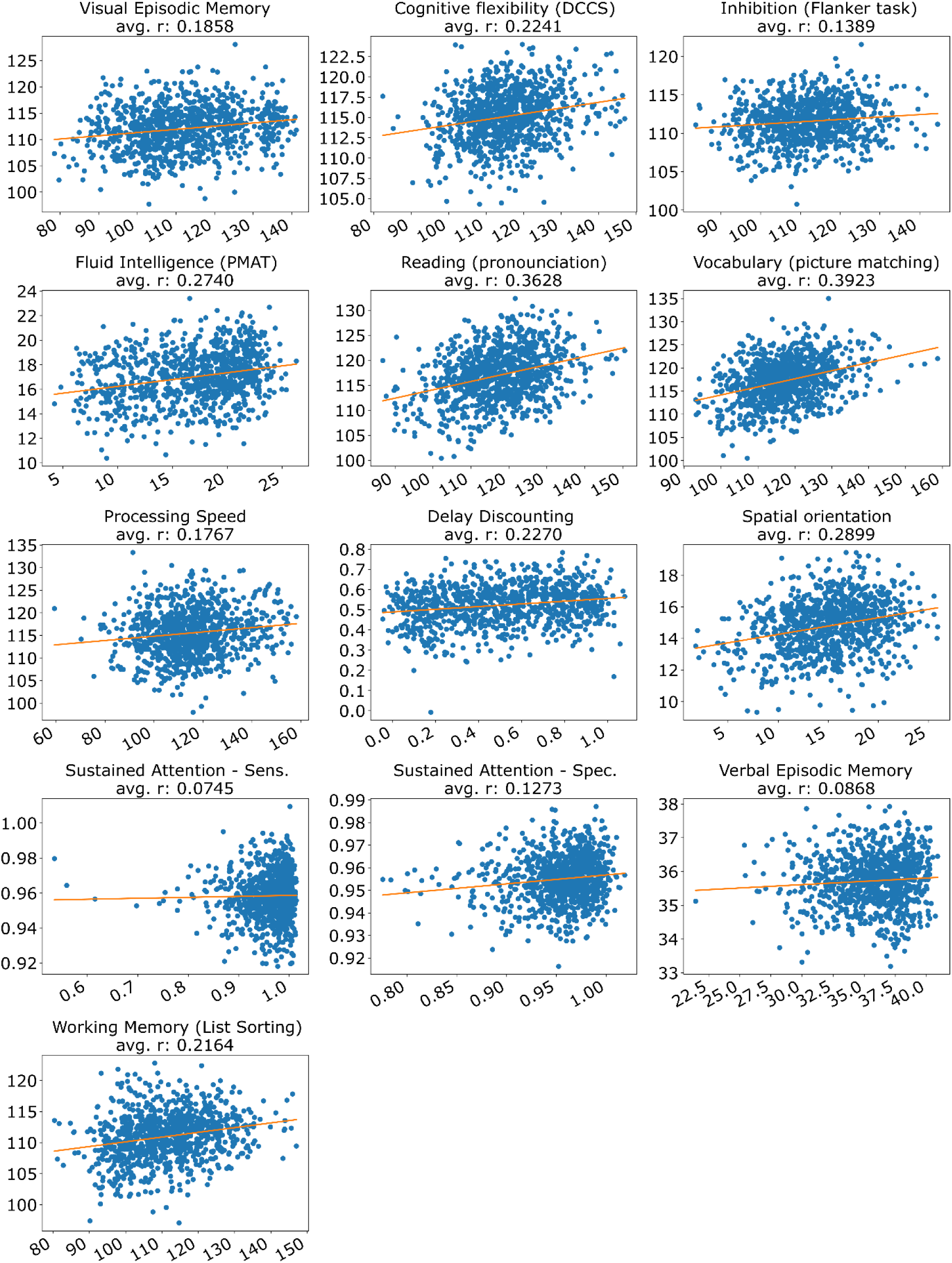
Scatterplots of predicted (BrainNetCNN) versus actual values for a curated set of 13 HCP cognitive measures for all 953 subjects. X-axis is the actual value. Y-axis is the predicted value. Each dot represents one subject. The orange line shows the best fit line obtained by least squares regression. The “avg. r” in each subplot title is the average Pearson’s correlation coefficient across 20 test folds, which was also reported in Figure 5.

**Figure S7.**
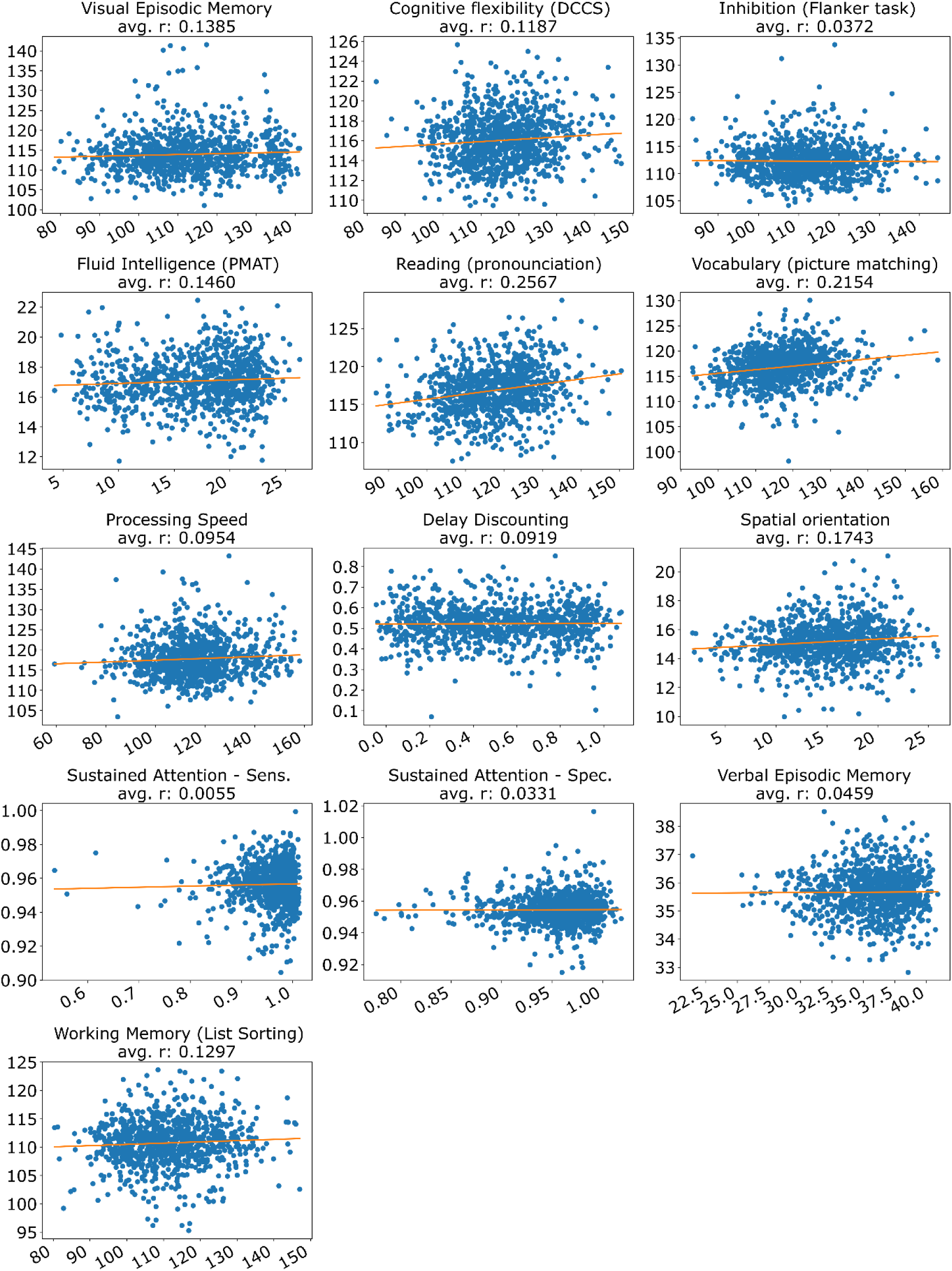
Scatterplots of predicted (GCNN) versus actual values for a curated set of 13 HCP cognitive measures for all 953 subjects. X-axis is the actual value. Y-axis is the predicted value. Each dot represents one subject. The orange line shows the best fit line obtained by least squares regression. The “avg. r” in each subplot title is the average Pearson’s correlation coefficient across 20 test folds, which was also reported in Figure 5.

**Figure S8.**
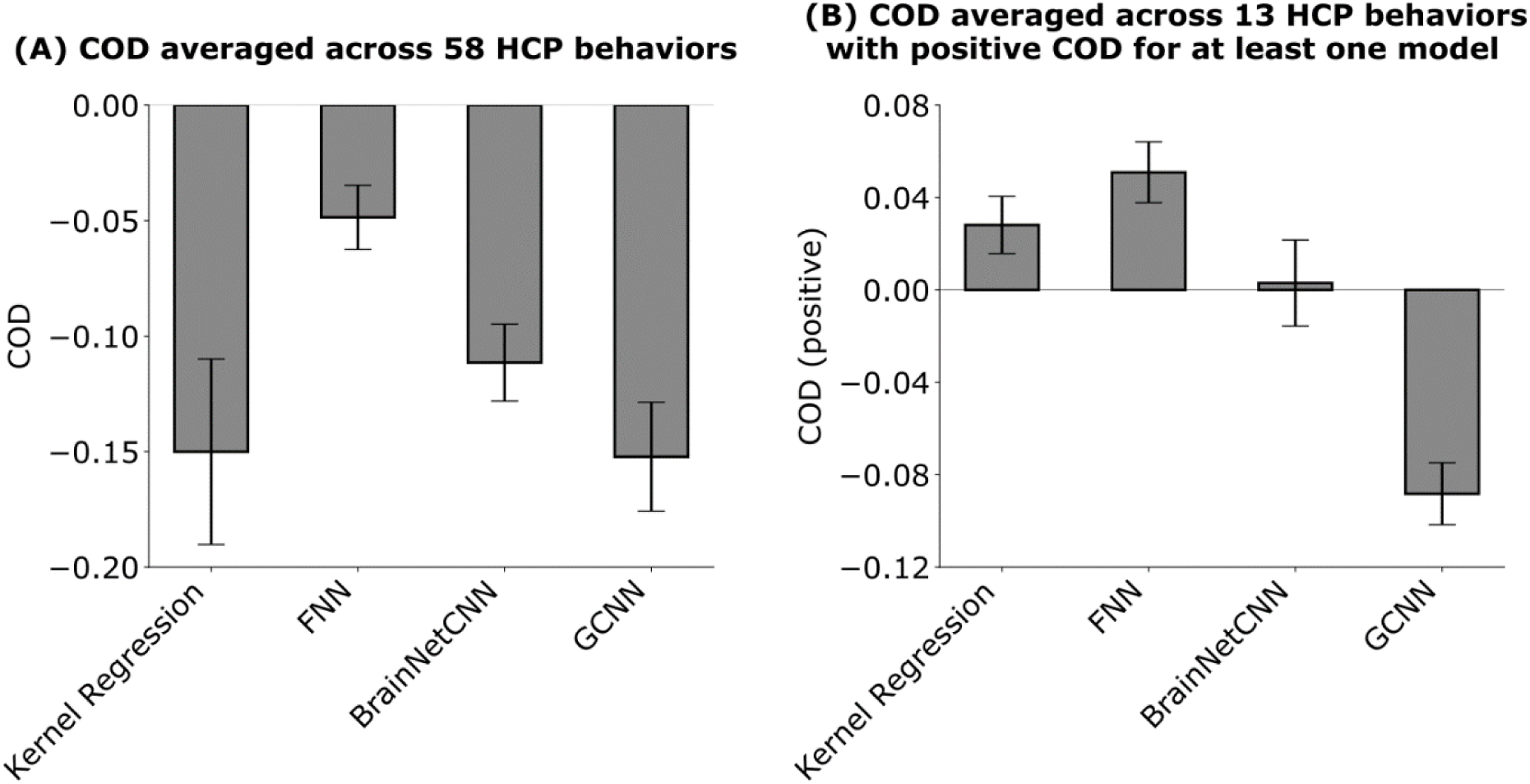
Prediction COD (coefficient of determination) averaged across HCP behavioral measures and 20 test folds. (A) Prediction COD averaged across all 58 behavioral measures. COD was computed for each test fold and each behavior and then averaged across the 58 behaviors and 20 test folds. Bars show the mean across test folds. Error bars show standard error of model performance across cross-validation folds. There was no statistical difference between kernel regression and all DNNs after correcting for multiple comparisons. (B) Prediction COD averaged across 13 behavioral measures. The negative COD in (A) suggests that there were a number of measures that were predicted poorly across all methods. Therefore, we considered a subset of measures with positive COD for at least one regression model (kernel regression, FNN, BrainNetCNN and GCNN). There were 13 such measures, so we averaged the COD across the 13 measures. Bars show the mean across test folds. Error bars show standard error of model performance across test folds.

**Figure S9.**
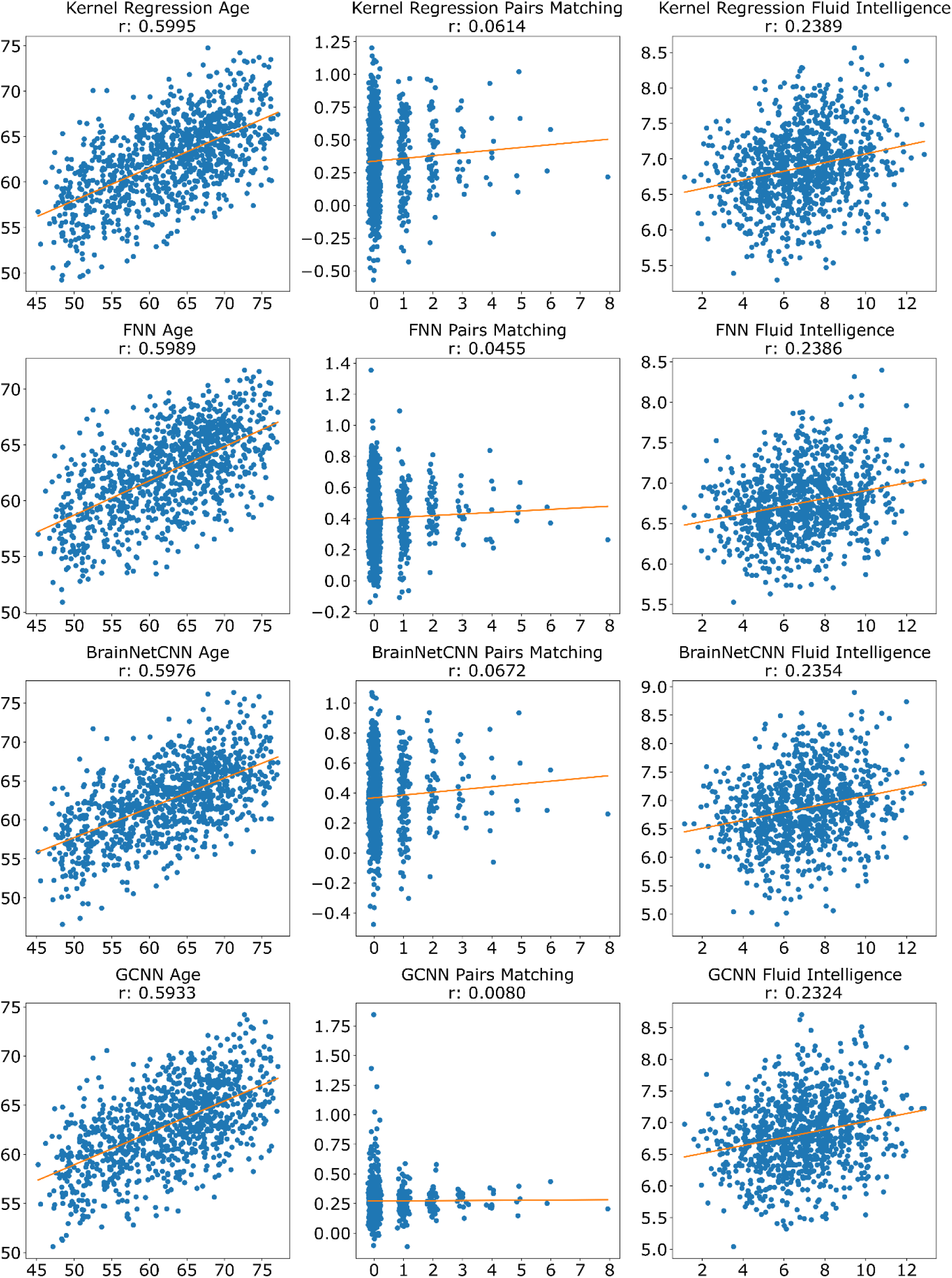
Scatterplots of predicted versus actual values for age, pairs matching and fluid intelligence for the 1000 UK Biobank subjects in the test set. X-axis is the actual value. Y-axis is the predicted value. Each dot represents one subject. The orange line shows the best fit line obtained by least squares regression. The “r” in each subplot title is the Pearson’s correlation coefficient between the predicted and actual values, which was also reported in Table 1.

**Figure S10.**
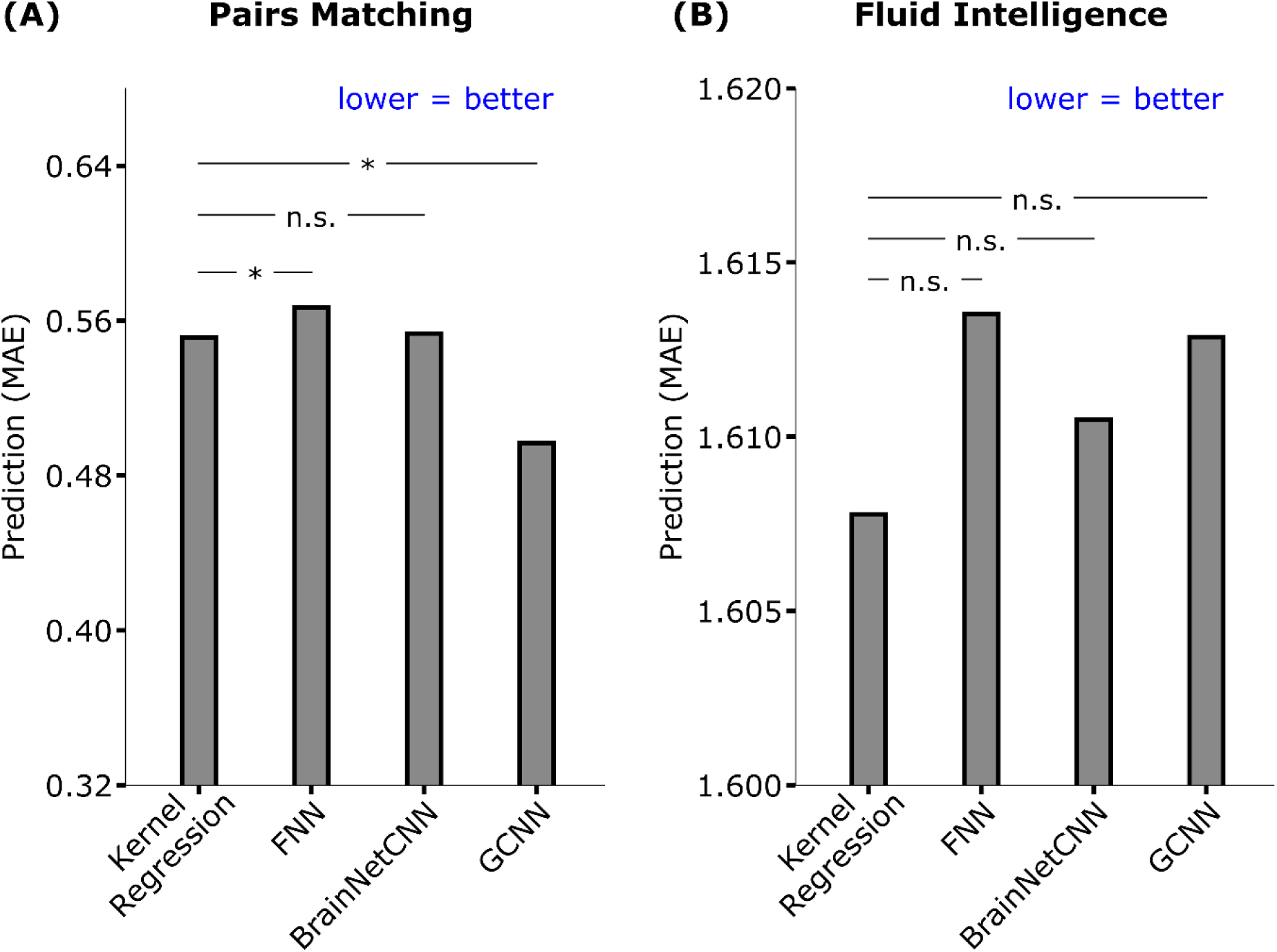
Prediction MAE of pairs matching and fluid intelligence in the UK Biobank. Lower values imply better performance. The horizontal lines represent statistical tests between kernel regression and the DNNs. “n.s” stands for not significant. “*” implies statistical significance after FDR (q < 0.05) correction. In the case of pairs matching, kernel regression was statistically better than GCNN (p = 1e-10), but statistically worse than BrainNetCNN (p = 0.009). However, we note that all algorithms performed worse than simply predicting the median pairs matching value in the training set, which would have yielded an MAE of 0.4.

**Table S2.** Prediction COD (Coefficient of determination) of age, pairs matching, and fluid intelligence in the UK Biobank. Higher values imply better performance. **Bold** indicates the best performance, although it does not imply statistical significance. We note that the negative values for pairs matching suggests that all four models were worse than simply predicting the mean for pairs matching. This was not the case for age or fluid intelligence.

| Model | Age | Pairs matching | Fluid intelligence |
| --- | --- | --- | --- |
| Kernel regression | **0.359** | -0.0751 | **0.0568** |
| FNN | 0.351 | -0.0339 | 0.0511 |
| BrainNetCNN | 0.355 | -0.0406 | 0.0506 |
| GCNN | 0.344 | **-0.0309** | 0.0512 |

**Figure S11.**
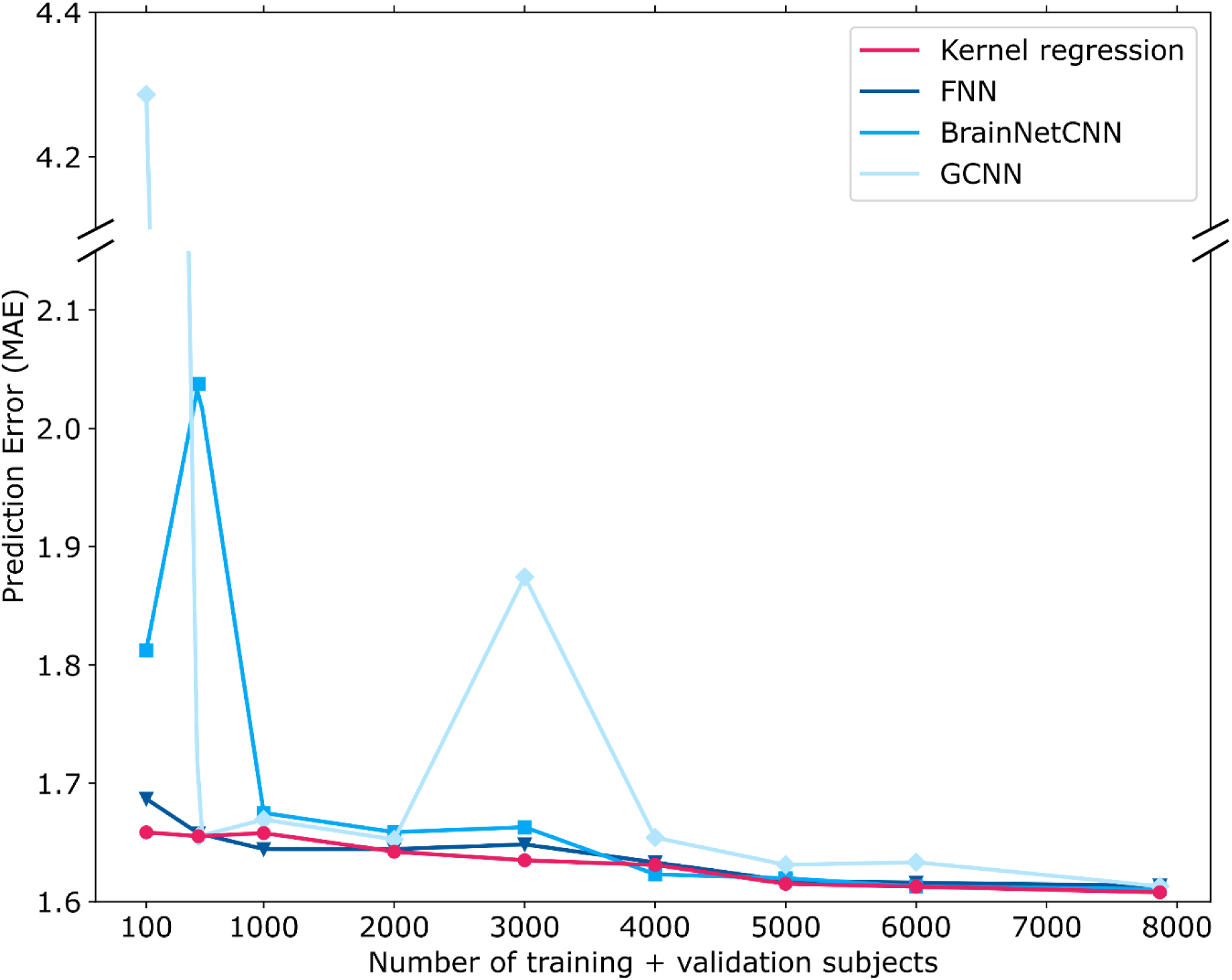
Prediction error (MAE) of fluid intelligence in the UK Biobank dataset with different number of training and validation subjects. Lower MAE represents better performance. The performance of all algorithms generally improved as the number of training and validation subjects increased. In the case of 100, 500, 1000 and 2000 subjects, 3/4 of the subjects were used for training and 1/4 of the subjects were used for validation. In the remaining cases, 1000 subjects were used for validation, while the remaining subjects were used for training. For all cases, test set comprised the same set of 1000 subjects. Kernel regression was highly competitive across all sample sizes.

**Table S3.** Prediction accuracy in the HCP dataset (Pearson’s correlation, averaged across 58 HCP behavioral measures and 20 test folds) for FNN (FC90net), BrainNetCNN and GCNN using reference hyperparameters (Kawahara et al., 2017; Parisot et al., 2017, 2018) versus our tuned hyperparameters (Figure 4). Results using our hyperparameters compared favorably with the results using the reference hyperparameters.

| Network | Performance using reference hyperparameters | Performance with our hyperparameters |
| --- | --- | --- |
| FNN | 0.119 | 0.121 |
| BrainNetCNN | 0.109 | 0.114 |
| GCNN | 0.0143 | 0.072 |

**Table S4.** Prediction accuracy in the UK Biobank for FNN (FC90net), BrainNetCNN and GCNN using reference hyperparameters (Kawahara et al., 2017; Parisot et al., 2017, 2018) versus our tuned hyperparameters (Figure 7). Results using our hyperparameters compared favorably with the results using the reference hyperparameters (except for GCNN and pairs matching).

| Model | Sex | Age | Pairs<br>matching | Fluid<br>intelligence |
| --- | --- | --- | --- | --- |
|  | Accuracy | Correlation | Correlation | Correlation |
| FNN (reference) | 0.895 | 0.581 | 0.032 | 0.231 |
| BrainNetCNN (reference) | 0.911 | 0.575 | 0.033 | 0.232 |
| GCNN (reference) | 0.855 | 0.446 | 0.051 | 0.122 |
| FNN (ours) | 0.916 | 0.599 | 0.045 | 0.239 |
| BrainNetCNN (ours) | 0.914 | 0.598 | 0.067 | 0.235 |
| GCNN (ours) | 0.916 | 0.593 | 0.008 | 0.232 |

**Table S5.** Test set prediction accuracy of the original GCNN implementation (provided in the GitHub repository of Kipf and Welling, 2017; https://github.com/tkipf/keras-gcn) and our implementation. Results were obtained using the toy data and hyperparameters provided by the GCNN GitHub repository (Kipf and Welling, 2017).

| GCNN | Accuracy |
| --- | --- |
| Original Implementation | 0.776 |
| Our Implementation | 0.817 |

**Table S6.** Test set prediction accuracy of the original BrainNetCNN implementation (provided in the GitHub repository of Kawahara et al., 2017; https://github.com/jeremykawahara/ann4brains) and our PyTorch and Keras implementations. Results were obtained using the toy data and hyperparameters provided by the BrainNetCNN GitHub repository (Kawahara et al., 2017).

| BrainNetCNN | Class 0 Corr | Class 0 MAE | Class 1 Corr | Class 1 MAE |
| --- | --- | --- | --- | --- |
| Original (100 epochs) | 0.043 | 0.230 | 0.439 | 0.229 |
| Original (1000 epochs) | 0.436 | 0.213 | 0.504 | 0.214 |
| PyTorch (100 epochs) | 0.062 | 0.231 | 0.072 | 0.235 |
| PyTorch (1000 epochs) | 0.720 | 0.155 | 0.266 | 0.240 |
| PyTorch (2000 epochs) | 0.720 | 0.155 | 0.258 | 0.233 |
| PyTorch (4000 epochs) | 0.765 | 0.186 | 0.531 | 0.244 |
| PyTorch (8000 epochs) | 0.821 | 0.154 | 0.737 | 0.226 |
| Keras (100 epochs) | 0.694 | 0.186 | 0.687 | 0.185 |
| Keras (1000 epochs) | 0.818 | 0.157 | 0.816 | 0.159 |

**Figure S12.**
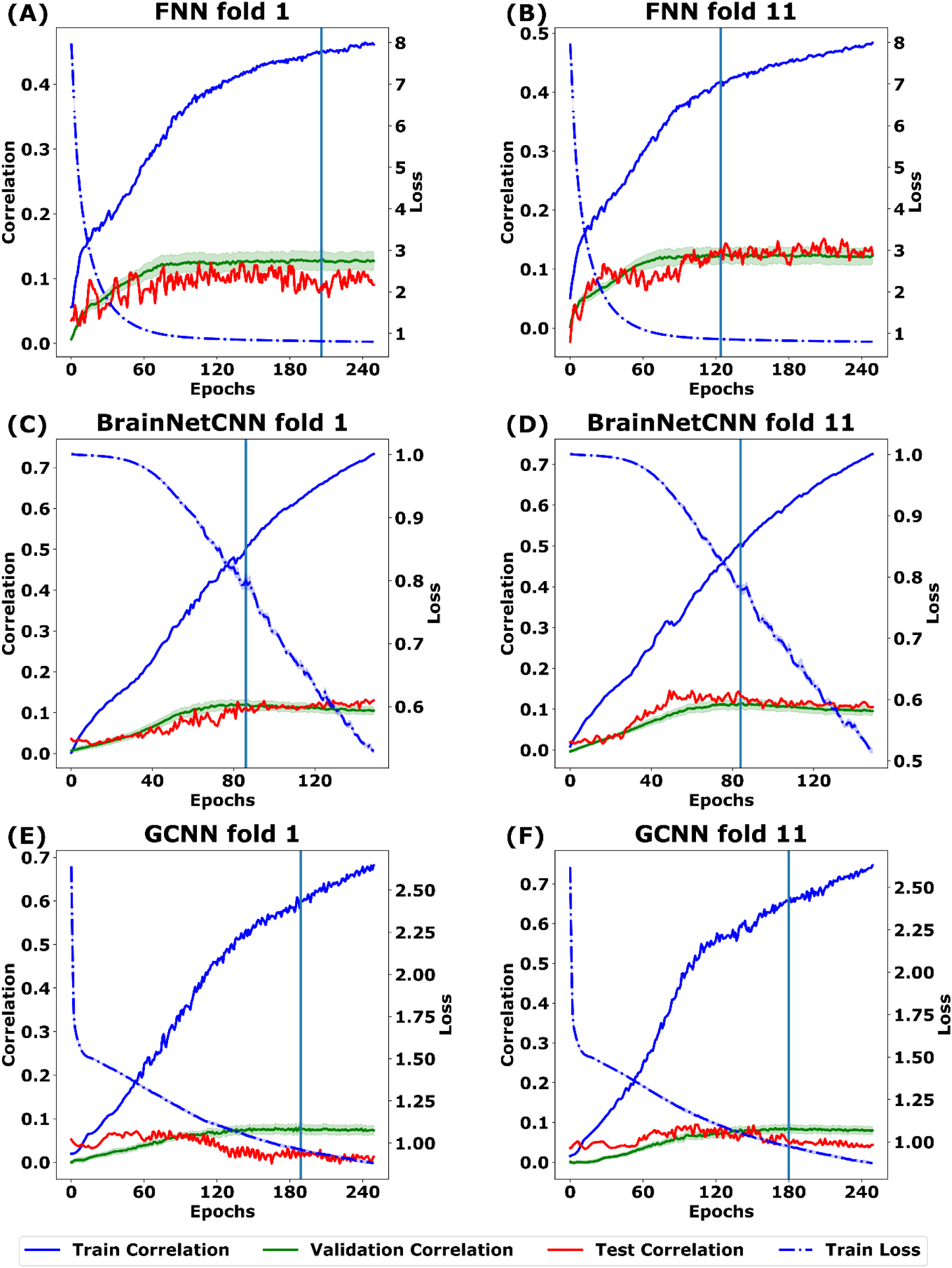
Learning curves of DNNs for folds 1 and 11 of the HCP dataset using the final set of hyperparameters. Blue dot curve shows the training loss averaged across 19 training curves during the inner-loop cross-validation. The green curve shows the validation correlation averaged across 58 behavioral measures and 19 validation curves during the inner-loop cross-validation. The shadow (for training loss and validation correlation) represents the standard error across 19 training and validation curves. The blue vertical line indicates the stopping point, which corresponded to the maximum validation correlation. The blue solid curve shows the training correlation when training on the 19 training folds (using the final set of hyperparameters), e.g., when fold 1 was the test fold, then folds 2 to 20 are the training folds. The red curve shows the testing correlation on the test fold.

**Figure S13.**
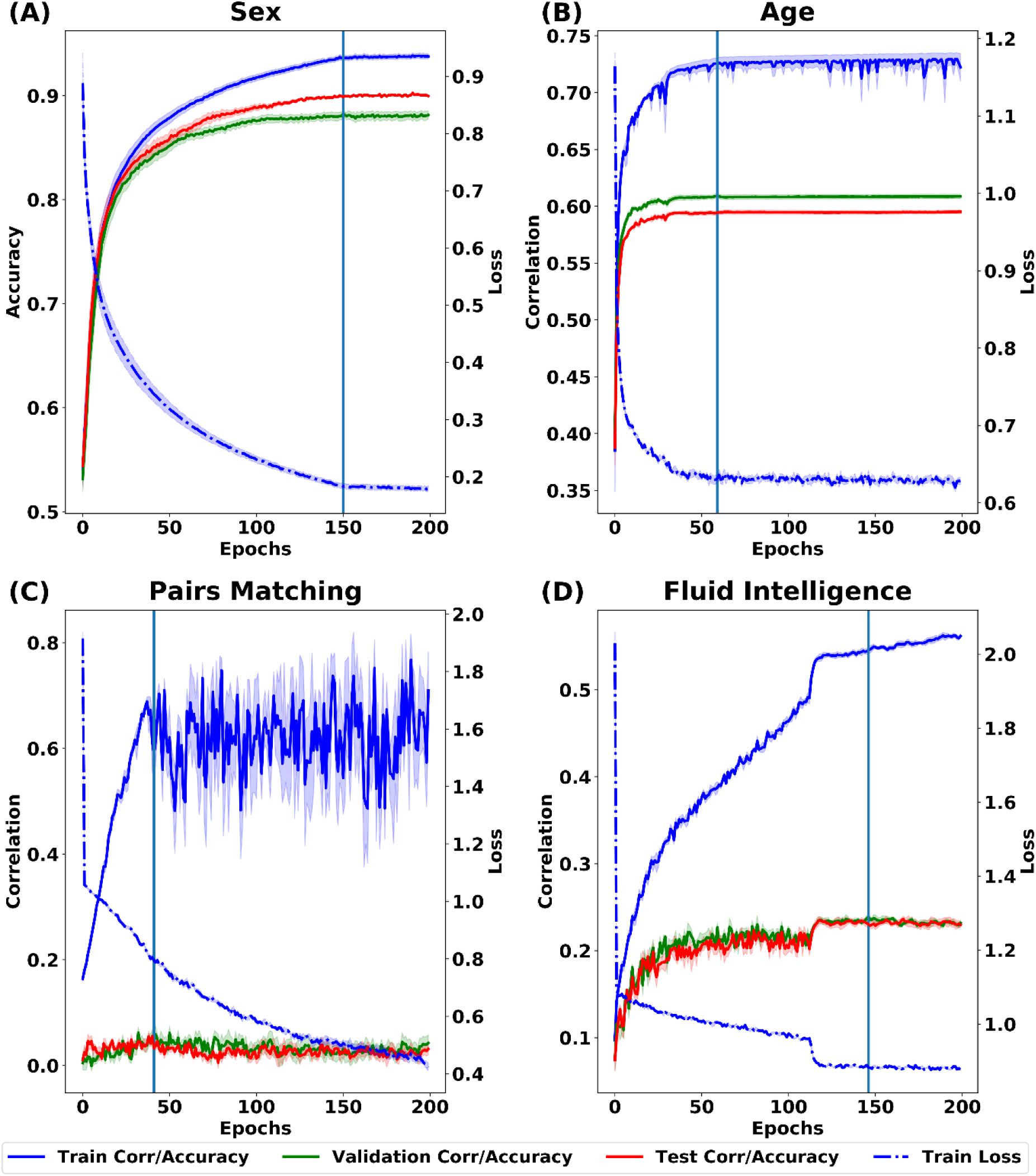
Learning curves of FNN for the UK Biobank using the final set of hyperparameters. Blue solid curve shows the training correlation (or accuracy for sex prediction). Blue dot curve shows the training loss. The green curve shows the validation correlation (or accuracy for sex prediction). The red curve shows the testing correlation (or accuracy for sex prediction). The shadow represents the standard error across the five ensemble runs. The blue vertical line indicates the stopping point, which corresponded to the best validation correlation/accuracy.

**Figure S14.**
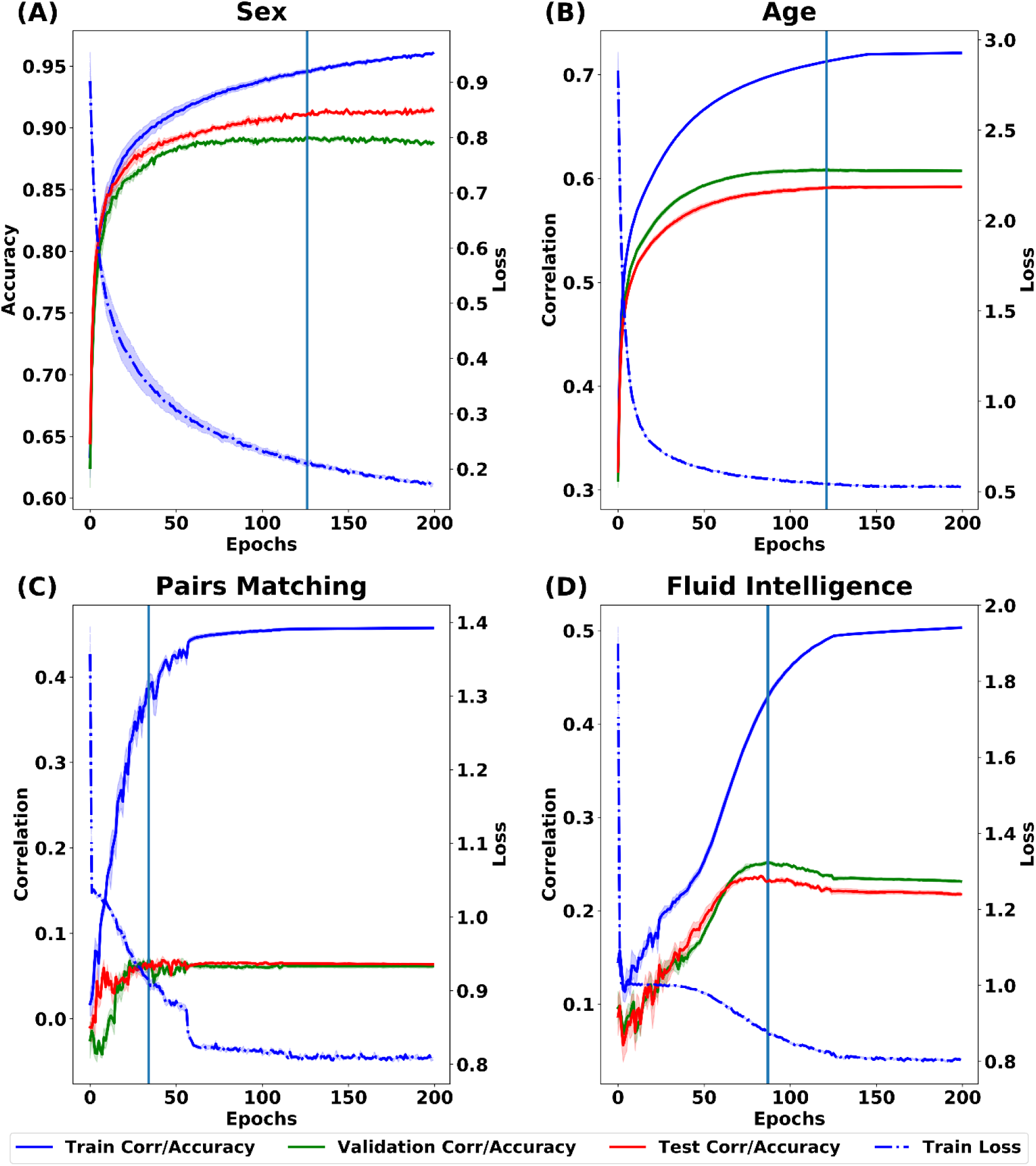
Learning curves of BrainNetCNN for the UK Biobank using the final set of hyperparameters. Blue solid curve shows the training correlation (or accuracy for sex prediction). Blue dot curve shows the training loss. The green curve shows the validation correlation (or accuracy for sex prediction). The red curve shows the testing correlation (or accuracy for sex prediction). The shadow represents the standard error across the five ensemble runs. The blue vertical line indicates the stopping point, which corresponded to the best validation correlation/accuracy.

**Figure S15.**
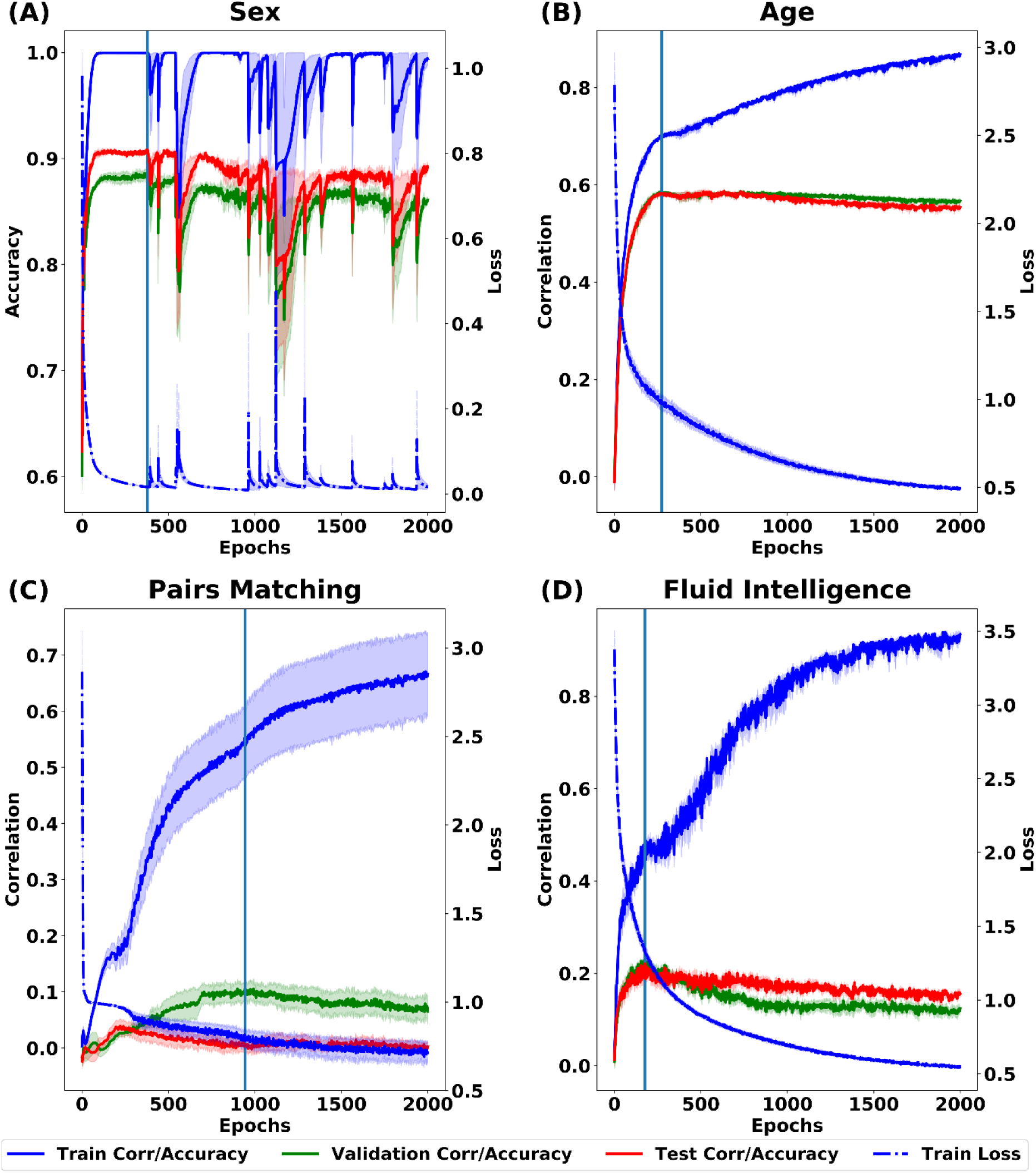
Learning curves of GCNN for the UK Biobank using the final set of hyperparameters. Blue solid curve shows the training correlation (or accuracy for sex prediction). Blue dot curve shows the training loss. The green curve shows the validation correlation (or accuracy for sex prediction). The red curve shows the testing correlation (or accuracy for sex prediction). The shadow represents the standard error across the five ensemble runs. The blue vertical line indicates the stopping point, which corresponded to the best validation correlation/accuracy.

1 The pairs matching task requires participants to memorize the positions of matching pairs of cards.

